# Estrogen receptor-related receptor (Esrra) induces ribosomal protein Rplp1-mediated adaptive hepatic translation during prolonged starvation

**DOI:** 10.1101/2024.01.09.574937

**Authors:** Madhulika Tripathi, Karine Gauthier, Reddemma Sandireddy, Jin Zhou, Priyanka Gupta, Suganya Sakthivel, Nah Jiemin, Kabilesh Arul, Keziah Tikno, Sung-Hee Park, Lijin Wang, Lena Ho, Vincent Giguere, Sujoy Ghosh, Donald P. McDonnell, Paul M. Yen, Brijesh K. Singh

**Author notes:** Authors share equal contribution (Madhulika Tripathi, Karine Gauthier). Corresponding author: Dr. Brijesh Kumar Singh, Cardiovascular and Metabolic Disorders Program, Duke– National University of Singapore (NUS) Medical School, Singapore 169857, Singapore.

## Abstract

Protein translation is an energy-intensive ribosome-driven process that is reduced during nutrient scarcity to conserve cellular resources. During prolonged starvation, cells selectively translate specific proteins to enhance their survival (adaptive translation); however, this process is poorly understood. Accordingly, we analyzed protein translation and mRNA transcription by multiple methods *in vitro* and *in vivo* to investigate adaptive hepatic translation during starvation. While acute starvation suppressed protein translation in general, proteomic analysis showed that prolonged starvation selectively induced translation of lysosome and autolysosome proteins. Significantly, the expression of the orphan nuclear receptor, estrogen-related receptor alpha (Esrra) increased during prolonged starvation and served as a master regulator of this adaptive translation by transcriptionally stimulating 60S acidic ribosomal protein P1 (Rplp1) gene expression. Overexpression or siRNA knockdown of Esrra expression *in vitro* or *in vivo* led to parallel changes in Rplp1 gene expression, lysosome/autophagy protein translation, and autophagy. Remarkably, we have found that Esrra had dual functions by not only regulating transcription but also controling adaptive translation via the Esrra/Rplp1/lysosome/autophagy pathway during prolonged starvation.

## 1. Introduction

Cells undergo adaptations to conserve energy during periods of nutrient scarcity or starvation. During acute starvation, there is suppression of protein translation, a vital cellular process by which ribosomes translate information from messenger RNA (mRNA) to synthesize proteins. However, during prolonged starvation, selective protein translation occurs in order to ensure cell survival (adaptive translation) ^1–3^.

Adaptive translation not only conserves vital resources but also shifts the utilization of cellular machinery and resources towards processes that help the cell to manage stress and enhance viability. It is tightly-controled and includes the selective protein translation of lysosome and autolysosome proteins involved in autophagy and other cellular catabolic processes ^4, 5^. In the liver, activation of lysosome-autophagy is crucial for maintaining the amino acid pool vital for translating proteins that oversee life-sustaining processes during starvation such as mitochondrial activity and β-oxidation of fatty acids ^6–11^. Although nuclear receptors are known to regulate transcription, their influence(s) on protein translation is not known. The orphan nuclear receptor, estrogen related receptor alpha (Esrra/ERRα) previously was shown to stimulate the transcription of mitochondrial genes ^10, 12^. Here, we show for the first time that prolonged starvation induces a novel Esrra-Rplp1 pathway to increase protein translation of lysosome and autolysosome proteins to sustain autophagy.

## 2. Results and Discussion

### 2.1 Selective protein translation activity was temporally regulated

Puromycin is an aminonucleoside that mimics the 3′ end of aminoacylated tRNAs and is incorporated into the C-terminus of nascent protein chains during the ribosome-mediated protein synthesis ^13^. To understand protein translation activity during acute and prolonged starvation, we performed puromycin-labeling of nascent proteins to evaluate their ribosome-mediated protein translation ^13–16^. Accordingly, mice were starved for 8, 24 or 48 h followed by puromycin-labeling (*i.p.* 20 mg/kg for 30 min just before the euthanization of mice) of newly translated proteins. The mice exhibited a time-dependent decrease in puromycin-labeled proteins at 8 h and 24 h in the liver followed by partial translational recovery at 48 h (**Figure 1A**). Similarly, primary human hepatocytes and mouse hepatic cells (AML12) cultured in serum-free media (serum starvation) for 0, 6, 24, 48, and 72 h followed by puromycin-labeling for a brief period (10 μg/ml for 15 min before the protein isolation), showed decreased puromycin-labeling from 6 h to 48 h serum starvation and partial recovery at 72 h in both hepatic cell lines (**Figures 1B,C**). We next performed polysome profiling to confirm the puromycin-labeled protein findings (**Figure 1D**). We observed decreased polysome peaks at 48 h serum starvation compared to 0 h suggesting reduced polysome-occupied mRNAs and translation activity at that time point. Interestingly, polysome peaks increased at 72 h serum starvation compared to 48 h, indicating there was a partial recovery in translation activity during prolonged starvation.

**Figure 1.**
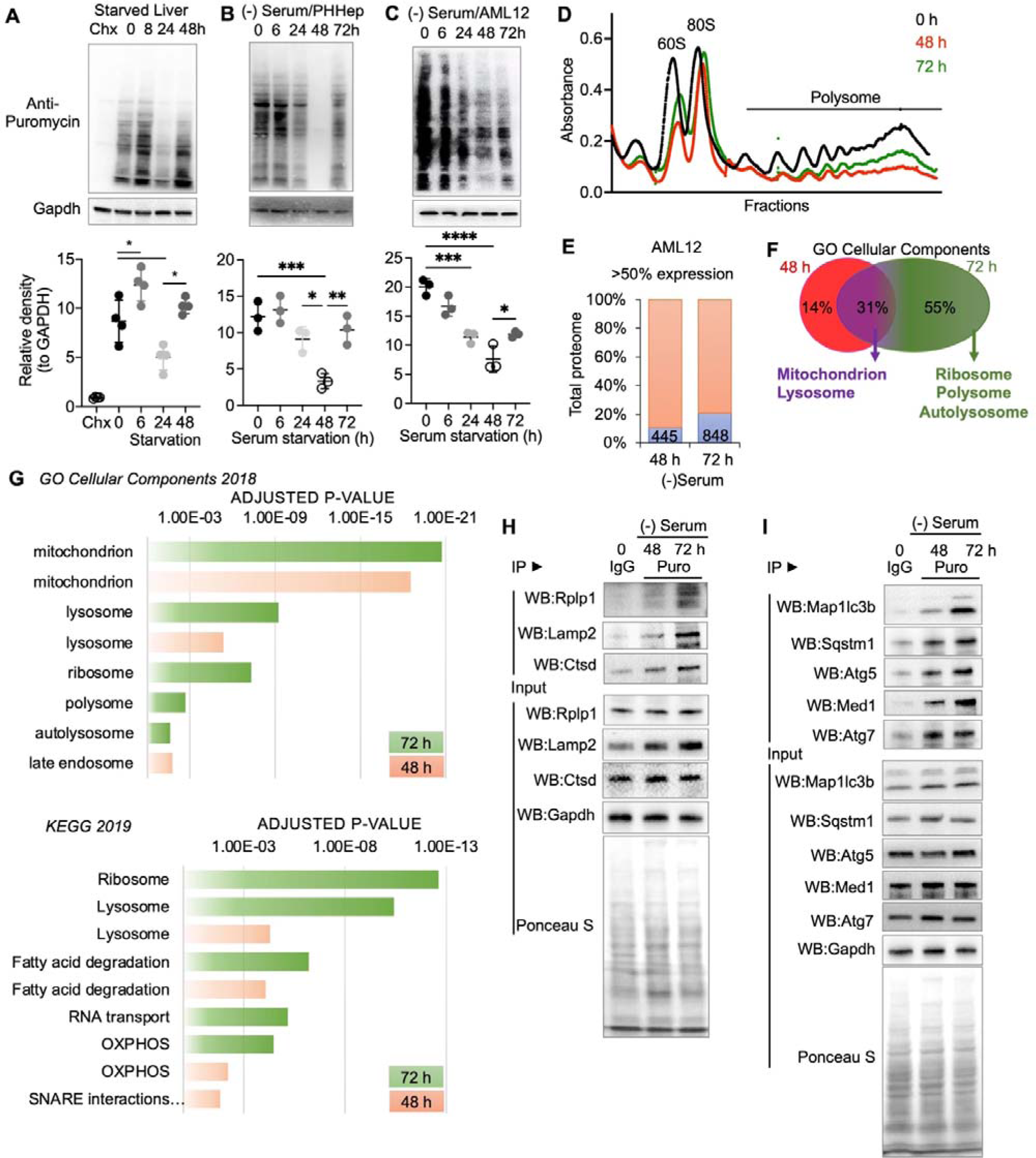
Hepatic protein translation was temporally regulated in starvation. (A-C) Representative Western blots of puromycin-labeled proteins in starved livers (n=4 per group), serum starved primary human hepatocytes (PHHep; n=3 per group), and AML12 cells (n=3 per group). Cycloheximide (Chx, 10 mg/kg body wight for 60 min) for translation inhibition and puromycin (Puro, 20 mg/kg body wight for 30 min) for protein labeling of nascent proteins were used just before euthanization for indicated time points. Dot plots below the Western blots represent relative density that is normalized to Gapdh. (D) Graph represents polysome profile in AML12 cells under 0, 48, and 72 h serum starvation, confirming puromycin-labeling results for translation activity. (E) Graph represents the percent of total number of upregulated (>50% expression compared to 0 h) proteins (absolute number in the bars) from a label-free quantification by mass spectrometry. (F) Venn analysis for common (purple) and exclusive (red for 48 h, and green for 72 h) pathways. (G) Graphs representing significant pathways analyzed using Gene Ontology (GO) Cellular Components and KEGG 2018 database on EnrichR platform (the Ma’ayan Lab, NY, USA). (H and I) Immunoprecipitation of puromycin-labeled proteins to analyze newly synthesized proteins and their detection using Western blotting. Ponceau S staining of the membranes showing protein content in each lane. Figures are showing a representative Western blot n=3. Levels of significance: *P<0.05; **P<0.01; ***P<0.001; ****P<0.0001.

To analyze translated proteins at 48 and 72 h serum starvation, we performed an unbiased label-free quantitative proteomic analysis in AML12 cells that underwent serum starvation for 0, 48 and 72 h (**Supplementary Figure 1A**). Surprisingly, despite the decrease in overall translation rate at 48 h serum starvation, approximately 10% of detectable proteins in the total proteome (445/4082 proteins) increased expression (> 50% higher expression than 0 h baseline) at 48 h, whereas 20% of detectable proteins in the total proteome (848/4066 proteins) increased at 72 h of serum starvation (**Figure 1E**), suggesting there was adaptive protein synthesis during prolonged starvation. We next analyzed all upregulated proteins at both time points to identify major induced pathways using the Gene Ontology Cellular Components (GO CC) 2018 databases and Kyoto Encyclopedia of Genes and Genomes (KEGG) 2019 (**Figure 1F, Supplementary Figures 1B**). Interestingly, 55% of the total up-regulated proteins were induced exclusively at 72 h of serum starvation and enriched for pathways regulating ribosomes, polysomes, RNA transport, ribosome biogenesis, autolysosomes, as well as lysosomes in the KEGG analysis (**Figure 1G, Supplementary Figure 1B, Supplementary Tables 1-3**). 14% of the total up-regulated proteins were exclusively induced at 48 h of starvation but did not correlate with any pathway (**Supplementary Table 1, 4 and 5**). Similar findings for pathways uniquely upregulated at 48 and 72 h of starvation also were observed in GO CC analyses (**Supplementary Tables 6-8**). Interestingly, 31% of the upregulated proteins were commonly induced at both 48 and 72 h of starvation (**Figure 1G)**. KEGG analyses of the commonly induced proteins revealed pathways related to lysosomes, fatty acid degradation, and mitochondrial oxidative phosphorylation (OXPHOS) (**Figure 1F and G, Supplementary Figure 1C, Supplementary Table 1**). Likewise, GO CC analysis confirmed that mitochondria and lysosomes were among the most significantly upregulated cellular components induced at both 48 and 72 h of starvation (**Supplementary Figure 1C; Supplementary Tables 6-8**).

### 2.2 Selective induction of ribosome, lysosome and autolysosome proteins in hepatic cells during prolonged starvation

When we analyzed the expression of individual proteins, we found that many of the proteins belonging to the ribosome pathway were downregulated after 48 h starvation compared to baseline but rose to higher levels than baseline after 72 h (**Supplementary Figure 1B**). Surprisingly, Rplp1, a ribosomal large subunit protein induced during embryonic development of the nervous system ^17^, was one of the most upregulated ribosomal proteins in adaptive translation occurring at 72 h serum starvation. Interestingly, the expression of many of the lysosomal proteins also were modestly upregulated at 48 h compared to the baseline (**Supplementary Figure 1B**) with the notable exceptions of Man2b1, Gla, and Gm2a which were downregulated. Significantly, the expression of almost all the lysosomal proteins, including Ctsd and Lamp2 as well as Man2b1, Gla, and Gm2a, were higher at 72 h than at 48 h (**Supplementary Figure 1B**). Additionally, expression of proteins involved in fatty acid oxidation (*e.g.,* Acox3, Acaa2, and Hadha) and OXPHOS (*e.g.,* Ndufa9, Ndufb9, and Sdha) modestly increased at 48 h and further increased at 72 h of starvation (**Supplementary Figures 1B**), consistent with the notion that sustained mitochondrial activity was required for fatty acid β-oxidation for cell survival during starvation. To validate the proteomics data, we performed puromycin immunoprecipitation of puromycin-labeled proteins at 0, 48 and 72 h and found that Rplp1 was highly enriched at 72 h whereas lysosomal proteins, Lamp2 and Ctsd were slightly enriched at 48 h, and further enriched at 72 h (**Figure 1H**). Moreover, qPCR analysis of polysome fractions from AML12 cells revealed that Rplp1, Lamp1, and Ctsd mRNA were enriched in polysome-occupied fractions (*i.e.,* mRNAs encoding high abundance proteins) at 72 h serum starvation compared to 0 or 48 h (**Supplementary Figures 1C and D**).

Starvation-induced autophagy is dependent upon lysosome-mediated catabolism ^18, 19^. Accordingly, we examined puromycin-labeling of several autophagy proteins: Map1lc3b, Sqstm1, Atg5, and Atg7 as well as Med1, an autophagy regulatory co-activator protein ^20^, and found they had similar temporal patterns of puromycin-labeling as lysosome proteins (**Figure 1I**) except for Atg7 which had increased puromycin-labeling at 48h but no further change at 72 h. Proteomic analysis also revealed that Map1lc3b, Sqstm1, Atg5, and Med1 protein expression increased at 72 h compared to 48 h whereas Atg7 protein expression remained the same at 72 h (**Supplementary Figure 1B**). Thus, activation of ribosome-mediated induction of protein translation helped maintain both lysosome and autophagy protein expression in hepatic cells at 72 h starvation.

### 2.3 Esrra regulation of ribosome-catalyzed translation of lysosome proteins in hepatic cells during prolonged starvation

We next performed a transcription factor analysis by Targeting Protein-Protein Interaction (TF PPIs) of upregulated proteins and identified Ppargc1a and Esrra as the transcription factors most likely involved in the increases in protein translation at 72 h of serum starvation (**Figure 2A**). Ppargc1a is a heterodimer partner of Esrra ^12^, so it is noteworthy that previous studies in *Ppargc1a* knockout (KO) mice displayed decreased Lamp2 and Ctsd expression in vascular smooth muscle cells ^21^. We and others also previously demonstrated Esrra transcriptionally regulated autophagy and mitophagy and stimulated its own transcription ^12, 22–24^. However, its role in the regulation of translation was not known.

**Figure 2.**
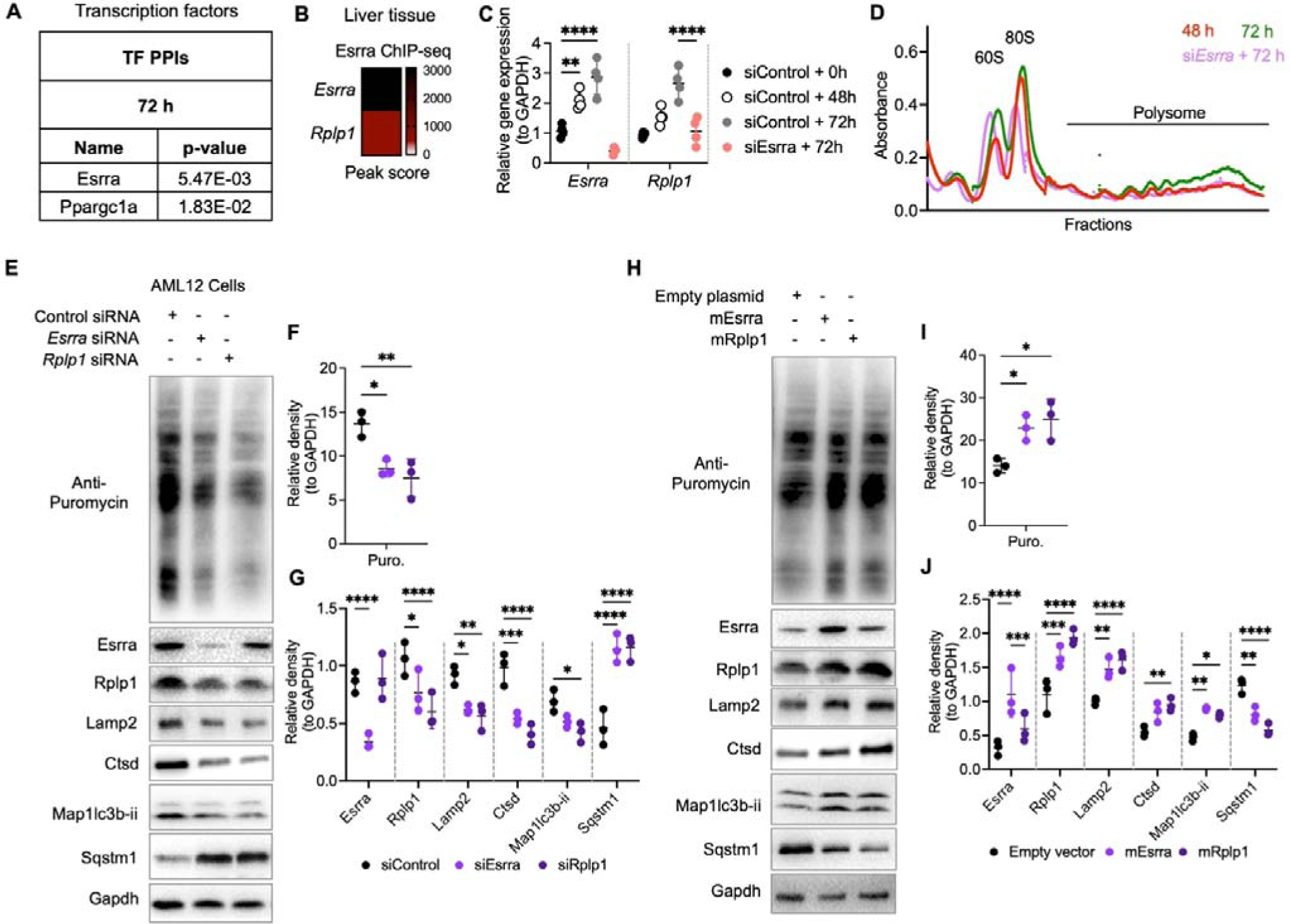
Esrra regulated ribosomal RPLP1 transcription, protein translation and the expression of lysosome and autophagy proteins in serum starvation. (A) Transcription factor analysis for >50% expressed proteins in 72 h serum starved AML12 cells compared to 48 h using Targeting Protein-Protein Interactions (PPIs) on EnrichR platform (the Ma’ayan Lab, NY, USA). (B) Heat map shows peak score of Esrra ChIP-seq analysis in liver tissues published previously ^12^. (C) RT-qPCR analysis of control and serum starved AML12 cells (for indicated time points) with or without *Esrra* siRNA. (D) Graph represents polysome profiling in AML12 cells under 48, 72 h and 72h serum starvation under si*Esrra* KD conditions. (E) Representative Western blots of AML12 cells treated with control, *Esrra* or *Rplp1* siRNA for 72 h. (F and G) Plots represent relative density of Western blots normalized to Gapdh. (H) Representative Western blots of AML12 cells treated with empty plasmid, Esrra or Rplp1 expressing plasmid for 72 h. (I and J) Plots represents relative density of Western blots normalized to Gapdh. Levels of significance: *P<0.05; **P<0.01; ***P<0.001; ****P<0.0001.

To understand the role of Esrra in the transcriptional regulation of *Esrra* and *Rplp1* genes during starvation, we analyzed hepatic Esrra ChIP-seq data and found that Esrra bound to its own promoter (**Figure 2B**), suggesting its transcription was involved in a positive-feedback loop ^12, 25^. Esrra also bound specifically to the *Rplp1* gene promoter (**Figure 2A**), the most upregulated ribosomal protein during serum starvation in AML12 cells (**Supplementary Figures 1B**). Of note, Esrra was not observed to bind to any of the other ribosomal gene promoters ranked among the top ten. Thus, to confirm if Esrra regulated Rplp1 transcription and Rplp1-mediated translation during serum starvation, we performed RT-qPCR analysis for *Esrra* and *Rplp1* gene expression (**Figure 2C**). Consistent with the hepatic Esrra ChIP-seq data, serum starvation induced Esrra and Rplp1 gene expression in a time-dependent manner from 48 h to 72 h serum starvation. Interestingly, *Esrra* siRNA knockdown (KD) in AML12 cells completely blocked the increase in Rplp1 gene expression at 72 h serum starvation and suggested Esrra regulated Rplp1 gene expression during prolonged serum starvation. Moreover, decreased polysome peaks in *Esrra* siRNA KD cells showed that *Esrra* KD completely inhibited protein translation at 72 h serum starvation (**Figure 2D**). Further, *Esrra* KD completely inhibited the enrichment of *Esrra*, *Rplp1*, *Lamp2*, and *Sqstm1* mRNA in polysome-occupied fractions (**Supplementary Figure 2A**); thus, supporting the notion that their active translation was Esrra-Rplp1-dependent. Additionally, *Esrra* or *Rplp1* KD in AML12 cells inhibited overall puromycin-labeled proteins and puromycin labeling of Lamp2, Ctsd, and Map1lc3b-ii proteins whereas puromycin labeling of Sqstm1 protein increased. These data suggested that reduced expression of either Esrra or Rplp1 proteins decreased protein translation, expression of lysosomal proteins, and autophagy (**Figures 2E-G**). In contrast, overexpression of either Esrra or Rplp1 in AML12 cells significantly increased puromycin-labeling of Lamp2, Ctsd, Map1lc3b-ii, and Sqstm1 proteins (**Figures 2H-J**), and suggested that activation or either Esrra or Rplp1 stimulated their translation.

Since autophagy is a highly adaptive and dynamic process where lysosomes play key roles in degradation and nutrient recovery during starvation ^26, 27^, we examined a more detailed time course of Esrra, Rplp1, lysosome protein (Lamp2 and Ctsd) and autophagy protein (Map1lc3b-ii and Sqstm1) expression in AML 12 cells at baseline, 6, 24, 48, and 72 h of serum starvation (**Supplementary** Figure 2B). Esrra expression increased during both acute and prolonged starvation whereas Map1lc3b-ii protein expression was high at 24 h, decreased at 48 h, and then increased again at 72h whereas Sqstm1 expression rose slightly at 48 h and returned back to baseline at 72, consistent with a decline and resumption of autophagy. These changes in Map1lc3b-ii protein expression were mirrored by expression of Rplp1, Lamp2, and Ctsd proteins (**Supplementary Figures 2B-D**) and further supported the role of Esrra and Rplp1 in regulating the recovery of both autophagy and lysosome proteins during prolonged starvation.

Next, we confirmed lysosomal activity, autophagy flux, and mitochondrial activity in AML12 cells transfected with or without Esrra siRNA during serum starvation as found earlier (**Figure 1F and G**). We used acridine orange (AO) staining to analyze lysosomal activity ^28^ and found, consistent with the pathway analysis data (**Figure 1G**), AO staining increased during serum starvation in control AML12 cells. In contrast, it decreased in Esrra siRNA KD cells (**Figures 3A and B**). These data confirmed that lysosomal activity increased during serum starvation and decreased when Esrra was knocked down. To demonstrate that Esrra-mediated autophagic flux increased during starvation, we transiently expressed RFP-eGFP-Map1lc3 plasmid in AML12 cells transfected with control and *Esrra* siRNAs, and undergoing serum starvation (**Figures 3C-E**). We observed increased red puncta formation in control siRNA treated cells (**Figure 3D**) indicating that there was an increase in autophagy flux since GFP fluorescence was quenched in active lysosomes ^29^. In contrast, there were increased yellow puncta in *Esrra* siRNA-treated cells in serum-containing media and even more in serum-free media (**Figure 3E**). The increased yellow puncta arose from increased fluorescence of both GFP and RFP due to inhibition of autophagy flux. We also used a lysosomal inhibitor, Bafilomycin A, to demonstrate starvation-induced autophagy flux ^12, 29^ as there was less accumulation of Map1lc3b-ii in *Esrra* KD cells than control cells (**Supplementary Figures 2E and F**), and more Map1lc3b-ii accumulation in Esrra-overexpressed cells than control cells (**Supplementary Figure 2G and H**).

**Figure 3.**
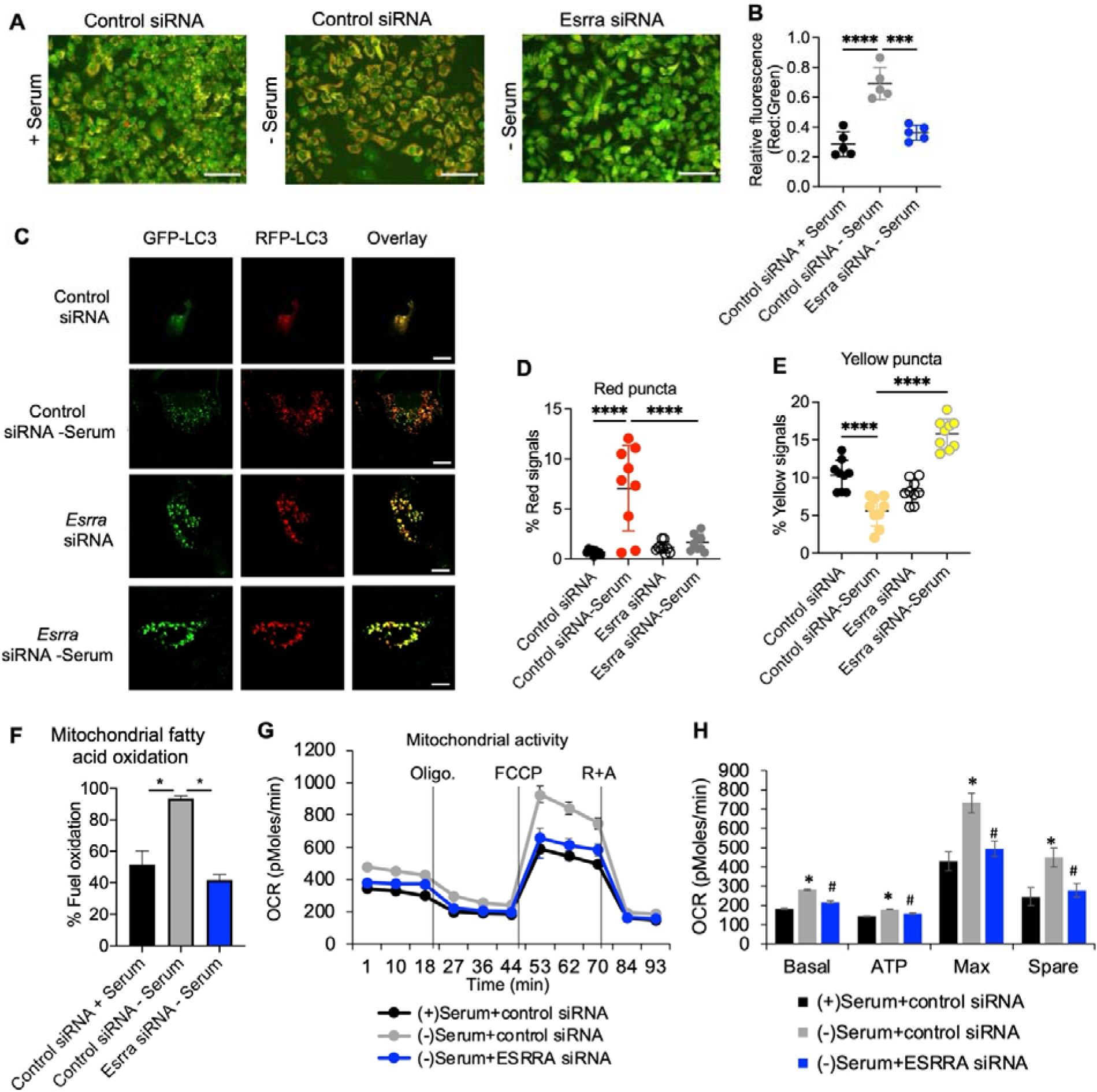
Esrra regulated lysosome activity, autophagy flux and mitochondrial activity during serum starvation. (A) Microscopic image showing acridine orange staining of AML12 cells treated with control siRNA or *Esrra* siRNA with or without 24 h serum starvation. Images are representative of three fields per group and three independent experiments. Scale bars, 200 μm. (B) Plot shows relative fluorescence that was measured using ImageJ (NIH, USA). n=5/groups. (C) Microscopic image showing RFP-GFP-LC3 expression in AML12 cells treated with control siRNA or *Esrra* siRNA with or without 24 h serum starvation. Images are representative of three fields per group and three independent experiments. Scale bars, 5 μm. (D and E) % red signals reflecting red puncta (D) for autolysosomes or % yellow signals reflecting yellow puncta (E) for autophagosomes. Colocalization analysis software CoLocalizer Pro 7.0.1 was used to analyze % red and yellow signals. (F) Mitochondrial fatty acid oxidation was analyzed in control siRNA and si*Esrra* treated AML12 cells using Seahorse XFe96 analyzer. (G and H) Mitochondrial oxidative phosphorylation (OXPHOS) as oxygen consumption rate (OCR) was analyzed in control siRNA and si*Esrra* treated AML12 cells using Seahorse XFe96 analyzer (G) and the parameters were calculated (H) as described in Methods section. Levels of significance: *P<0.05; **P<0.01; ***P<0.001; ****P<0.0001.

Esrra previously was shown to be a major regulator of mitochondrial biogenesis, β-oxidation of fatty acids, and mitophagy by inducing the expression of key genes involved in these processes ^10, 12^. In this connection, we found that *Esrra* siRNA KD inhibited mitochondrial fatty acid fuel oxidation, respiratory activity, OXPHOS, and ATP production in AML12 cells undergoing starvation (**Figure 3F-H**). These findings demonstrated that Esrra coordinately regulated autophagy and these key metabolic and energy production processes to enable an integrated response during starvation.

### 2.4 Esrra regulation of ribosome-catalyzed translation of lysosome proteins in hepatic cells prevented cell death during prolonged starvation

We next examined Esrra/Rplp1 regulation of lysosome and autophagy protein translation during prolonged starvation by puromycin-labeling proteins in cells grown in normal or serum free media either as basal or containing a highly specific Esrra inverse agonist, C29 ^25, 30–32^ (**Supplementary Figures 3A-C**). C29 decreased Esrra protein expression and prevented induction of Rplp1, reduced selective puromycin-labeling of proteins, and diminished Lamp2, Ctsd, and Map1lc3b-ii protein expression. Sqstm1 significantly accumulated due to reduced lysosome activity and late block autophagy. Furthermore, micrographs clearly showed that inhibition of Esrra by C29 during 72 h serum starvation induced cell death (**Supplementary Figure 3D**), suggesting that Esrra regulation of adaptive protein translation was critical for cell survival during prolonged serum starvation. To determine whether the increased autophagy and lysosome protein expression at 72 h serum starvation was due to translational recovery or enhanced transcription of these genes, we performed RT-qPCR analysis in these samples, and found that although *Esrra, Rplp1, Lamp2, Ctsd, Map1lc3b,* and *Sqstm1* gene expression significantly increased at 72h serum starvation, *Ctsd, Map1lc3b,* and *Sqstm1* gene expression increased even when Esrra was inhibited by C29 (**Supplementary Figure 3E**). Moreover, we also confirmed that inhibiting Esrra using C29 during serum starvation in primary human hepatocytes decreased puromycin-labeling of proteins and induced significant cell death (**Supplementary Figure 3F and G**). These findings were discordant with the protein expression data and demonstrated that the increased expression of lysosomal and autophagy proteins at 72 h serum starvation was primarily due to Esrra-mediated Rplp1-dependent translation rather than increased transcription. The induction of *Ctsd, Map1lc3b,* and *Sqstm1* gene expression during prolonged starvation, even after C29 treatment, suggested there was alternative regulation by transcription factorss such as TFEB and FOXOs rather than Esrra^33, 34^. Taken together, our data clearly showed that continuous replenishment of autophagy proteins was required to maintain autophagy flux and cell survival during prolonged starvation. Further, we confirmed that Esrra regulation of autophagy flux included translation of lysosome proteins in addition to the transcriptional mechanisms described previously ^10, 24, 35^.

### 2.5 Esrra, Rplp1, lysosome and autophagy proteins in mice were co-regulated during fasting and refeeding

To better understand Esrra regulation of ribosome-dependent translation during starvation and its effects on lysosome-autophagy function *in vivo*, we analyzed liver tissues from mice fasted for 24h and then refed mice for 6 h (**Figures 4A and B**). Consistent with our *in vitro* data, protein expression of Esrra, Rplp1, Lamp2, Ctsd, and Map1lc3b-ii increased significantly, whereas Sqstm1 expression significantly decreased during prolonged fasting suggesting there was activation of the Esrra-Rplp1-lysosome pathway and increased autophagy flux during prolonged fasting. Interestingly, upon refeeding there was significantly reduced hepatic expression of Esrra, Rplp1, Lamp2, Ctsd, and Map1lc3b-ii expression whereas Sqstm1 was accumulated when compred to fed. These data strongly suggested reduced Esrra/Rplp1-mediated adaptive translation led to decreased lysosomal activity and a late block in autophagy in refed mice ^12, 36^.

**Figure 4.**
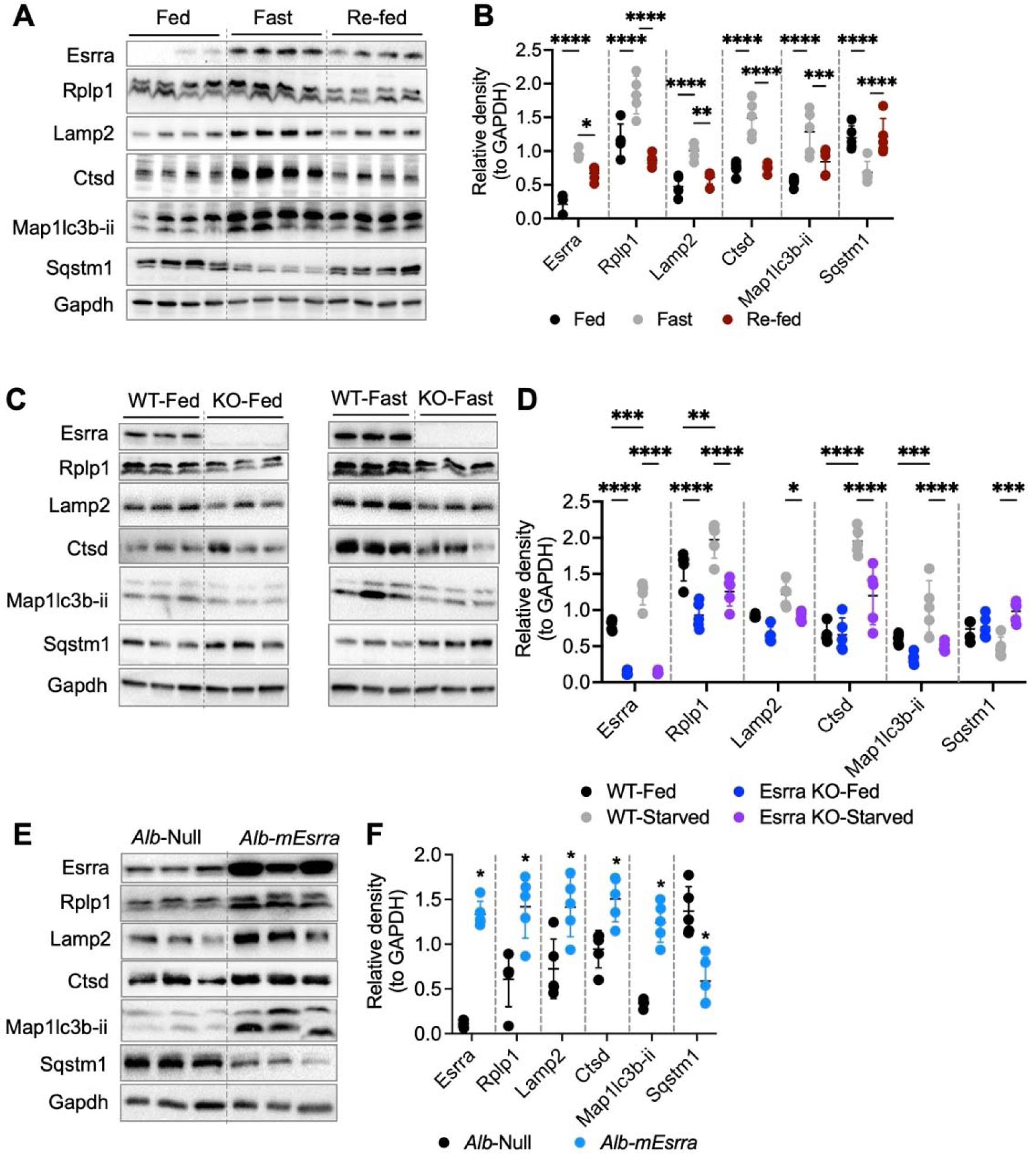
Esrra regulated temporal changes in Rplp1-lysosome axis and autophagy in mice livers. (A) Representative Western blots of fed, 24 h fasted and 6h refed livers. (B) Plots represent relative density of corresponding Western blots normalized to Gapdh (n=5 per group). (C) Representative Western blots of fed and 24 h starved livers from WT and *Esrra* KO mice. (D) Plots represent relative density of corresponding Western blots normalized to Gapdh (n=5 per group). (E) Representative Western blots of livers from liver-specific overexpressed Esrra (*Alb-mEsrra*) or control (*Alb*-null) mice. (F) Plots represent relative density of corresponding Western blots normalized to Gapdh (n=5 per group). Levels of significance: *P<0.05; **P<0.01; ***P<0.001; ****P<0.0001.

To demonstrate that Esrra/Rplp1-mediated adaptive translation was essential for autophagy *in vivo,* we examined the effects of inhibiting *Esrra* in mice treated with the inverse agonist of Esrra, XCT790 ^12, 37, 38^. XCT790-treated mice had significantly decreased hepatic expression of Esrra, Rplp1, Lamp2, Ctsd, Map1lc3b-ii and increased Sqstm1 (**Supplementary Figures 4A and B**). Electron micrographs also confirmed that hepatic lysosome/autolysosome number was significantly decreased in XCT790-treated mice compared to vehicle-treated controls, suggesting that Esrra regulated lysosome synthesis (**Supplementary Figures 4C and D**). Furthermore, we compared wild-type (WT) and *Esrra* knockout (KO) mice in the fed state and after 24h starvation (**Figures 4C and D**). In the fed state, *Esrra* KO mice showed a significant decrease in Rplp1 expression and a downward trend in the expression of Lamp2 and Map1lc3b-ii proteins compared to WT mice. Fasting increased Esrra, Rplp1, Lamp2, Ctsd, and Map1lc3b-ii protein expression, and had no significant effect on Sqstm1 protein in WT mice. In *Esrra* KO mice, these increases in the protein expression during starvation were significantly attenuated, and Sqstm1 expression increased during starvation suggesting that there was a late block in autophagy when Esrra was not expressed (**Figures 4C and D**). Likewise, hepatic *Esrra* overexpression in mice increased the hepatic expression of Rplp1, Lamp2, and Ctsd proteins and promoted autophagy flux as evidenced by the increased Map1lc3b-ii and decreased Sqstm1 expression (**Figures 4E and F**).

To better understand Esrra’s effects on hepatic protein translation during starvation *in vivo*, we injected mice with vehicle (control) or C29 during starvation, and measured protein puromycin labeling at baseline, 8, 24, and 48 h starvation. Hepatic protein translation decreased from 8 to 24 h and then significantly increased at 48 h compared to 24 h in control mice suggesting there was some recovery at 48h (**Figures 5A and B**). In contrast, there was a significant inhibition of overall puromycin labeling from 8 to 48 h starvation in C29-injected mice compared to their fed state (0 h). Esrra, Lamp2, Ctsd, and Map1lc3b-ii protein translation progressively increased from 8 h to 48 h starvation in vehicle-treated mice (**Figures 5A, C-F).**

**Figure 5.**
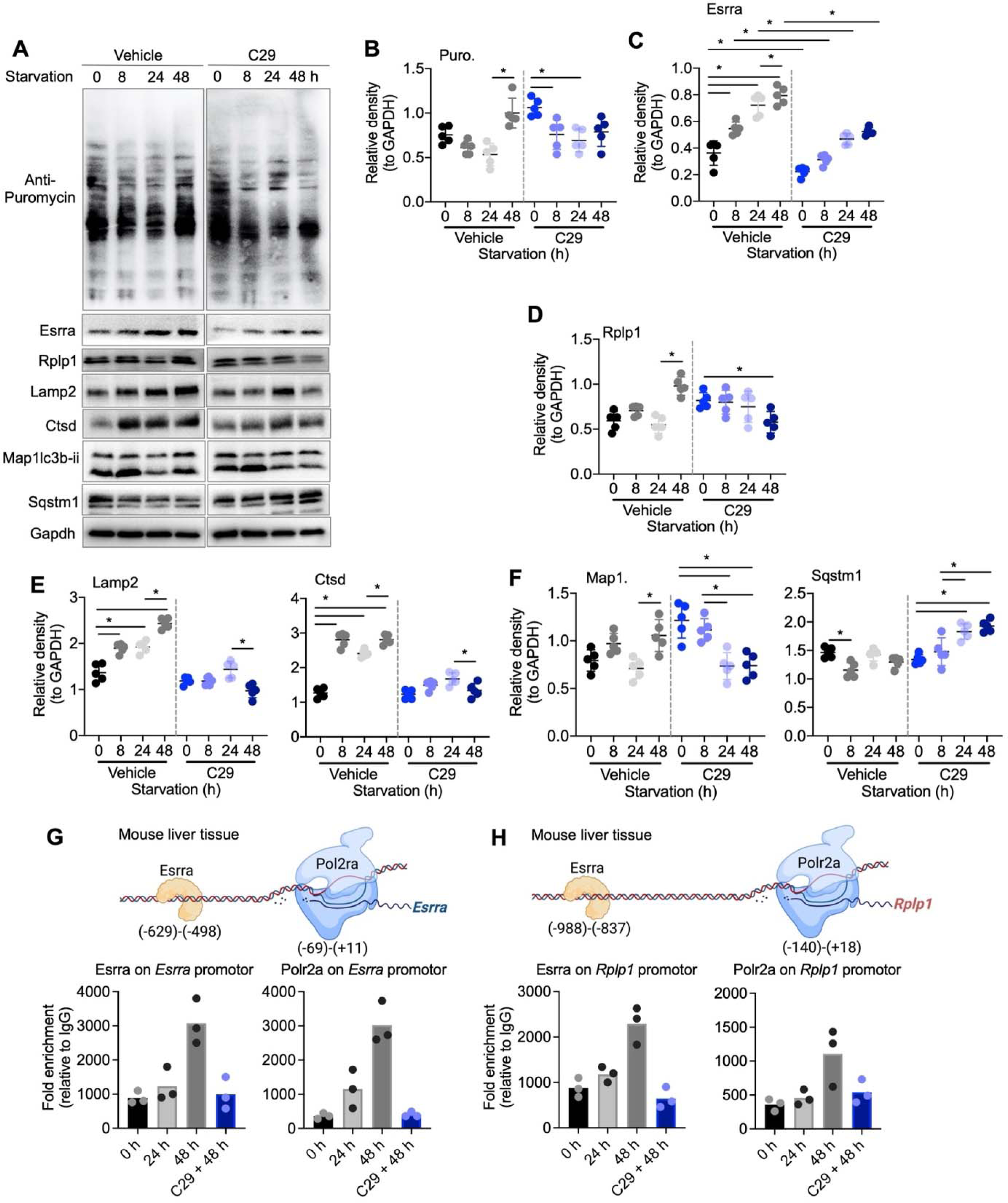
Esrra regulated temporal changes in Rplp1-lysosome axis and autophagy in mice livers. (A) Representative Western blots of starved mice livers at indicated time points treated with vehicle or Esrra inhibitor C29 (n=5 per group). (B-F) Plots represent relative density of corresponding Western blots normalized to Gapdh.J(G and H) ChIP-qPCR analysis of Esrra and Polr2a binding on *Esrra* or *Rplp1* promoters respectively. Experiment was performed three times independently. Levels of significance: *P<0.05.

However, Esrra, Rplp1, Lamp2, and Ctsd protein translation was significantly lower in C29-treated mice than vehicle-treated mice at each time point. Map1lc3b-ii protein translation also was inhibited and Sqstm1’s was increased in C29-treated mice starved for 24 h and 48 h (**Figures 5A, C-F**).

### 2.5 Esrra and Polr2a binding to the Esrra and Rplp1 promoters were necessary for autophagy-mediated β-oxidation of fatty acids

We identified strong Esrra response element (ESRRE) binding sites on the *Esrra* and *Rplp1* promoters using previously published mouse liver Esrra ChIP-seq databases ^12, 39^ (**Figure 2B**). We then used chromatin immunoprecipitation-qPCR (ChIP-qPCR) assays to observe Esrra and Polr2a binding on *Esrra* and *Rplp1* gene promoters at these ERREs and the TATA-box in the livers from mice starved for 24h and 48 h treated with and without C29 (**Figure 5G and H**). Both Esrra and Polr2a increased their binding to *Esrra* and *Rplp1* promoters in untreated mice at 48 h starvation while C29 completely inhibited Esrra and Polr2a binding on *Esrra* and *Rplp1* gene promoters at this time point (**Figure 5G and H**). C29 treatment also significantly inhibited hepatic *Esrra* and *Rplp1* mRNA expression in mice starved for 48 h (**Supplementary Figure 4E**). Interestingly, these mice nevertheless had higher *Lamp2, Ctsd, Map1lc3b* and *Sqstm1* mRNA expression than control mice, despite lower protein translation than controls suggesting alternative transcriptional mechanisms when Esrra was inhibited (**Supplementary Figure 4E**). On the other hand, the progressive increases in *Esrra* and *Rplp1* mRNAs correlated with increased protein translations during starvation in control mice. Furthermore, serum β-hydroxybutyrate (β-HB), a marker of hepatic fatty acid β-oxidation regulated by lipophagy ^12, 40^, was significantly higher in control mice after 24 starvation compared to 0 h, and further increased after 48 h starvation (**Supplementary Figure 4F**). In contrast, β-HB did not further increase in C29-injected mice starved for 48h. These data showed that Esrra-dependent Rplp1-mediated translation of lysosome-autophagy proteins was required for sustained fatty acid β-oxidation during starvation *in vivo*.

In summary, our comprehensive study examined the intricate mechanisms by which cells adapted to nutrient scarcity; particularly, the role of adaptive protein translation for cell survival during prolonged starvation. This investigation highlighted the dynamic nature of adaptive protein translation during starvation, revealing a temporal pattern that involved initial suppression of protein translation followed by selective translation of lysosome and autophagy proteins to maintain cellular homeostasis and survival during prolonged starvation.

One of the important findings of this study was the identification of Esrra as a key regulator of adaptive protein translation. We showed there was a temporal and selective regulation of lysosome and autophagy protein translation during prolonged starvation that was orchestrated by Esrra-mediated induction of *Rplp1* gene expression (See **Supplementary Figure 4G**). Thus, Esrra had dual functions of not only increasing *Rplp1* and *Esrra* gene expression, but also stimulating the translation of lysosome and autophagy proteins to increase autophagy, amino acid availability, and lysosome and mitochondrial protein translation, as well as to improve their functions. It is possible that agents increasing Esrra expression or mimicking its activity potentially may enhance Esrra-Rlplp1-lysosome protein translation. Such compounds, by functioning as “fasting” analogs, could be useful therapeutically to increase hepatic fatty acid β-oxidation in metabolic disorders such as NASH. It is however noteworthy that the induction of Esrra and the inhibition of mTOR activity also may have synergistic effects during fasting ^25, 41^.

Interestingly, Esrra enhanced autophagy in other cell types ^10, 12, 22–24, 42^ so this pathway may be involved in tissues other than the liver. Currently, the role(s) of Esrra-Rplp1-lysosome protein translation on autophagy in other hepatic, metabolic, or malignant conditions is not known. In particular, it remains to be determined whether the decreased Esrra expression observed in metabolic diseases such as NASH, obesity, and diabetes cause aberrant starvation response, autophagy, or protein translation that could be restored by induction of the Esrra-Rplp1-lysosome protein translation pathway.

## 4. Methods

### Animal studies

#### Fed-Fast-Refed mice

8-10 weeks old male C57BL/6J mice (n=5/group) mice were kept in empty cages (without husk bedding and NCD) for 24 (Fast group), and mice from Fast group were refed for 6 h with NCD (Refed). Mice fed with normal chow control diet (NCD) were considered as Fed group.

#### Starvation studies

8-10 weeks old male C57BL/6J mice (n=5/group) mice were kept in empty cages (without husk bedding) for 8, 24, 48 h. C29 (10 mg/kg body weight) was injected *i.p.* for the chronic inhibition of Esrra ^25^. Cycloheximide (Chx; 10 mg/kg body weight) was injected *i.p.* 60 min before euthanization, while puromycin (Puro, 20 mg/kg body weight) was injected *i.p.* 30 min before euthanization.

#### *Esrra* KO mice

*Esrra* KO mice are described elsewhere ^25, 43^. Esrra WT and *Esrra* KO mice (n=5/group) were starved for 24 h or left fed.

#### *Alb*-m*Esrra* overexpression

8 weeks old male C57BL/6J mice (n-5/group) were used for liver-specific *Esrra* overexpression (*Alb-mEsrra*). Mice were injected with AAV8-*Alb*-*mEsrra* (5X10^11^ gc/mice) via tail vein and housed for four weeks with no other intervention ^44^.

#### General mouse care and ethics statement

Mice were purchased from InVivos, Singapore and, housed in hanging polycarbonate cages under a 12 h/12 h light/dark schedule at Duke-NUS vivarium. Mice were simple randomized before grouping and fed different diets and normal water, or fructose treated *ad libitum*. Animals were euthanized in CO2 chambers. All mice were maintained according to the Guide for the Care and Use of Laboratory Animals (NIH publication no. One.0.0. Revised 2011), and the experiments performed were approved by the IACUCs at SingHealth (2015/SHS/1104) and (2020/SHS/1549).

### Cell cultures

#### AML12 cells

(ATCC® CRL-2254™) were cultured as indicated elsewhere ^45, 46^. Starvation medium (DMEM:F12 mix with Pen/Strep lacking serum, ITS and Dexamethasone) was used for serum starvation experiments. Puromycin (10 μg/ml) was added briefly for 15 min before harvest the cells ^13^ whereas C29 (5 μM) was added to media for indicated time to inhibit Esrra ^25^. Bafilomycin A1 (5 nM) was used to analyze autophagy flux ^12^.

#### Primary human hepatocytes

(5200, ScienCell) were cultured as indicated elsewhere ^44, 47^. Puromycin (10 μg/ml) was added for 15 min before harvest ^13^ whereas C29 (5 μM) was added to media for indicated time ^25^. Detailed methodology regarding the animal models and cell culture studies can be found in Supplementary Methods. Other methodological details for quantitative proteomics, *in vitro* polysome profiling and gene manipulation, measurements serum β-hydroxybutyrate (β-HB/Ketone bodies), ChIP-qPCR analysis in liver tissues, RNA/protein expression, Seahorse OCR analysis, statistical analysis etc. can be found in the Supplementary Methods section of Supplementary Information file.

## Acknowledgements and funding details

The authors like to acknowledge that the research is funded by the Ministry of Health (MOH), and National Medical Research Council (NMRC), Singapore, grant number NMRC/OFYIRG/0002/2016 and MOH-000319 (MOH-OFIRG19may-0002), Duke/Duke-NUS Research Collaboration Pilot Project Award (Duke/Duke-NUS/RECA(Pilot)/2022/0060), and KBrFA (Duke-NUS-KBrFA/2023/0075) to BKS; NMRC/OFYIRG/077/2018 to MT; and CSAI19may-0002 to PMY; Duke-NUS Medical School and Estate of Tan Sri Khoo Teck Puat Khoo Pilot Award (Collaborative) Duke-NUS-KP(Coll)/2018/0007A to JZ. This work is also partially supported by grants from the Louisiana Clinical and Translational Science Center (NIGMS 2U54GM104940), the National Heart Lung and Blood Insititute, NIH, USA (NHLBI R01HL146462-01), and the Khoo Bridge Fund, Singapore (KBrFA/2022/0060) to SG. The illustrations were made on BioRender.com.

## Author’s contributions

MT, BKS, LW, SG, experimental design/execution, data analysis; KG, Esrra KO mice experiments; KT, RS, KA, wet lab work; JZ, fast-refed mice experiments and hepatic-ESRRA overexpression in mice; NJ, LH, polysome profiling; VG, generated ESRRA KO mice, finalized manuscript; SP, DPM, shared C29, experimental suggestions, finalized manuscript; YW, BHB, EM, finalized manuscript; MT, BKS, PMY, drafted/finalized manuscript, provided financial support.

