## Supplemental data and methods for "Estrogen receptor-related receptor (Esrra) induces ribosomal protein Rplp1-mediated adaptive hepatic translation during prolonged starvation"

**Supplementary material**

**Supplementary material contains:**

**Supplementary figures (1-4)**

**Supplementary methods**

**Supplementary tables (1-8)**

**Supplementary Figures**

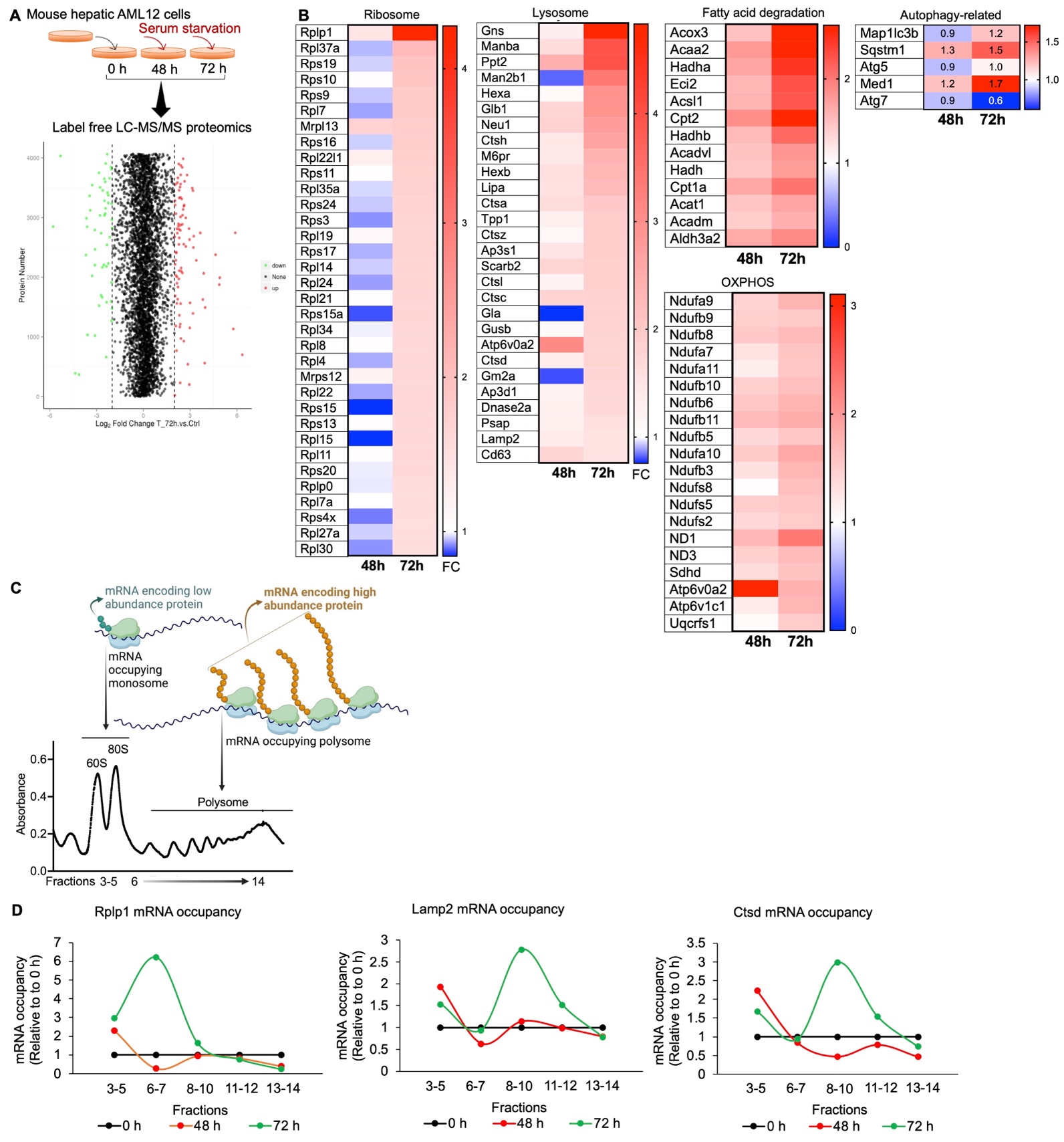

**Supplementary Fig. 1. Global protein translation was temporally regulated during serum starvation.** (A) Illustration shows serum starvation strategy and Label Free Quantitative (LFQ) proteomic identification in Volcano plot. (B) Heat maps representing the differential expression of proteins regulating Ribosome, Lysosome, Fatty acid oxidation, Oxidative Phosphorylation (OXPHOS), and Autophagy-related proteins in 48 and 72 h serum starved AML12 cells when compared to 0 h time point. (C) Illustration showing fractions 3-5 (from polysome profile) represents mRNA occupied by monosome, while fractions 6-14 represent mRNA occupied by polysomes (representing their active translation). (D) qPCR analysis of mRNA occupancy in monosome (fractions 3-5) or polysome (fractions 6-14) fractions.

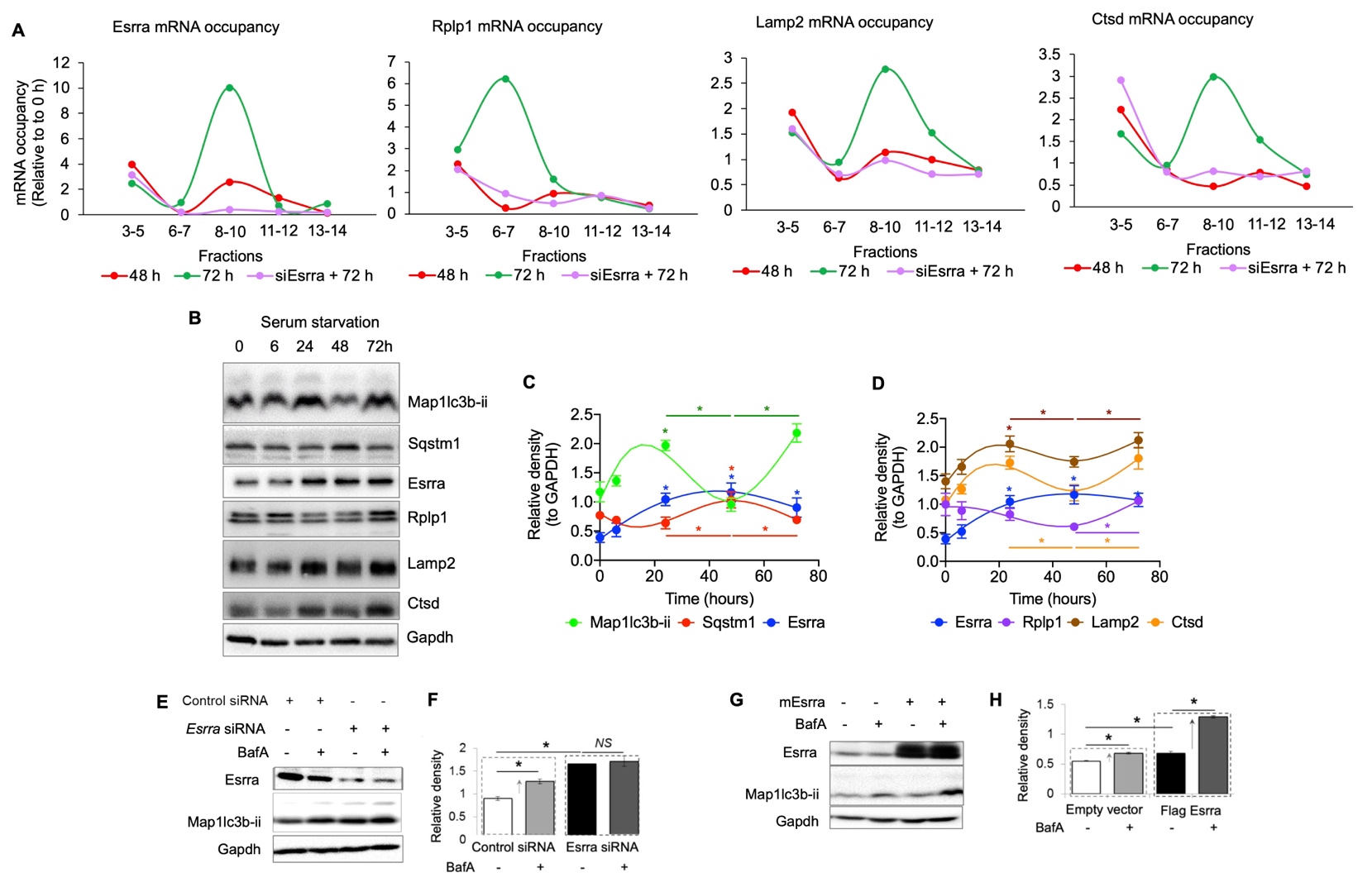

**Suppl. Fig. 2. Esrra regulated lysosome and autophagy activities under serum starvation.** (A) qPCR analysis of mRNA occupancy in monosome (fractions 3-5) or polysome (fractions 6-14) fractions. (B) Representative Western blots of serum starved AML12 cells for indicated time points (n=3 per group). (C and D) Plots represent relative density of corresponding Western blot normalized to Gapdh showing temporal changes in the protein expression. (E) Representative Western blots of AML12 cells treated control siRNA or *Esrra* siRNA with or without bafilimycin A1 (Baf) for 4 h. (F) Plot represents relative density of Western blots normalized to Gapdh. (G) Representative Western blots of AML12 cells treated empty plasmid or Esrra expressing plasmid with or without bafilimycin A1 (Baf) for 4 h. (H) Plot represents relative density of Western blots normalized to Gapdh.

Levels of significance: *P<0.05. *Compared with controls or as indicated, #compared with serum starved controls.

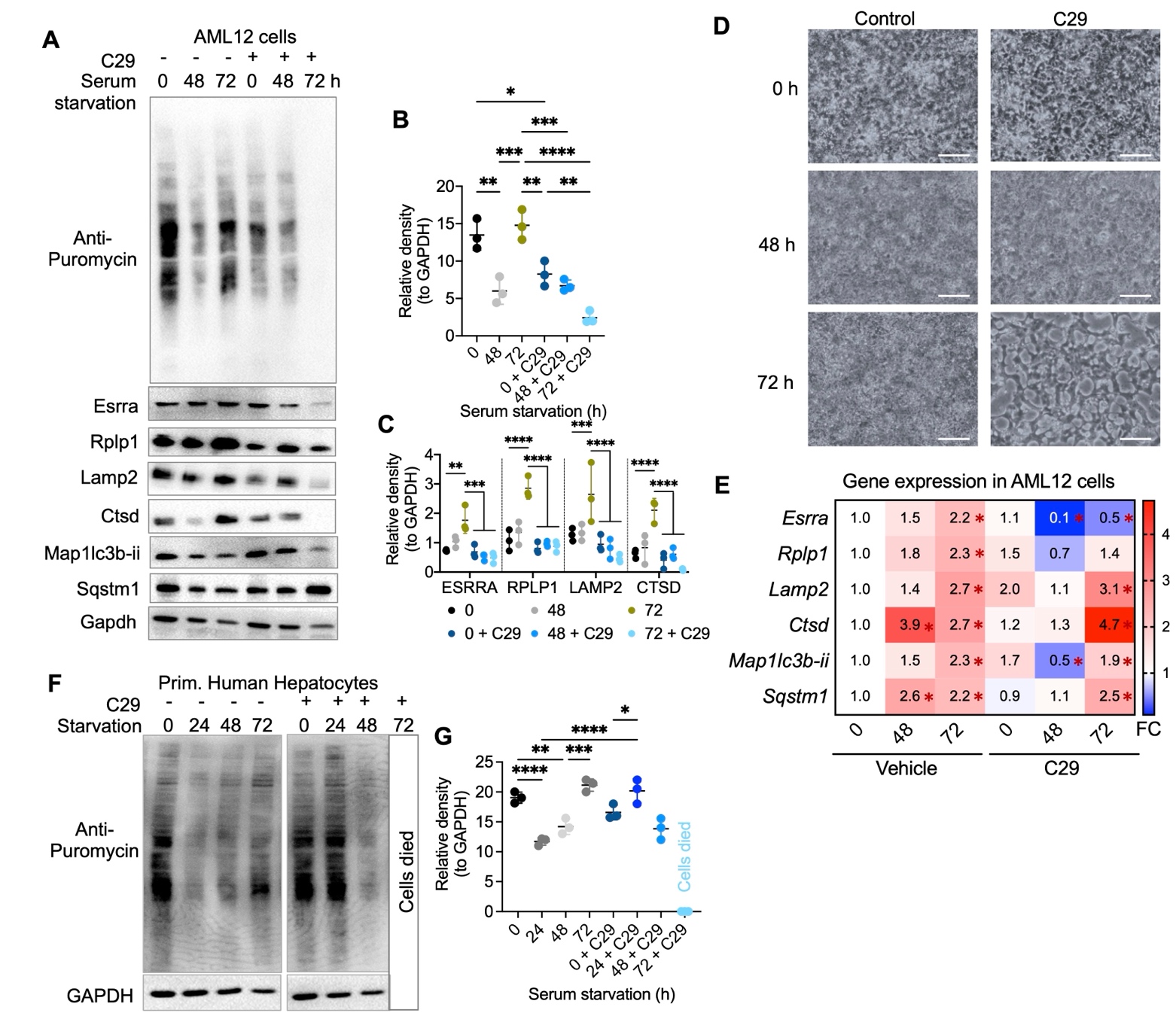

**Suppl. Fig. 3. Esrra regulated temporal changes in Rplp1-lysosome axis and autophagy *in vitro* during serum starvation.** (A) Representative Western blots of vehicle or C29 treated serum starved AML12 cells for indicated timepoints (n=3 per groups). (B and C) Plots represent relative density of corresponding Western blots normalized to Gapdh. (D) Microscopic image showing morphology of serum starved AML12 cells for 0, 48, 72 h and 72 h with C29. C29 treated 72 h starved cells showed massive cell death. Scale bars, 200 μm. (E) RT-qPCR analysis of relative gene expression in AML12 cells for indicated timepoints (n=3 per groups). Gene expression was normalized to *Gapdh*. (F) Representative Western blots of vehicle or C29 treated serum starved primary human hepatocytes for indicated timepoints (n=3 per groups). (G) Plots represent relative density of corresponding Western blots normalized to Gapdh.

Levels of significance: *P<0.05; **P<0.01; ***P<0.001; ****P<0.0001.

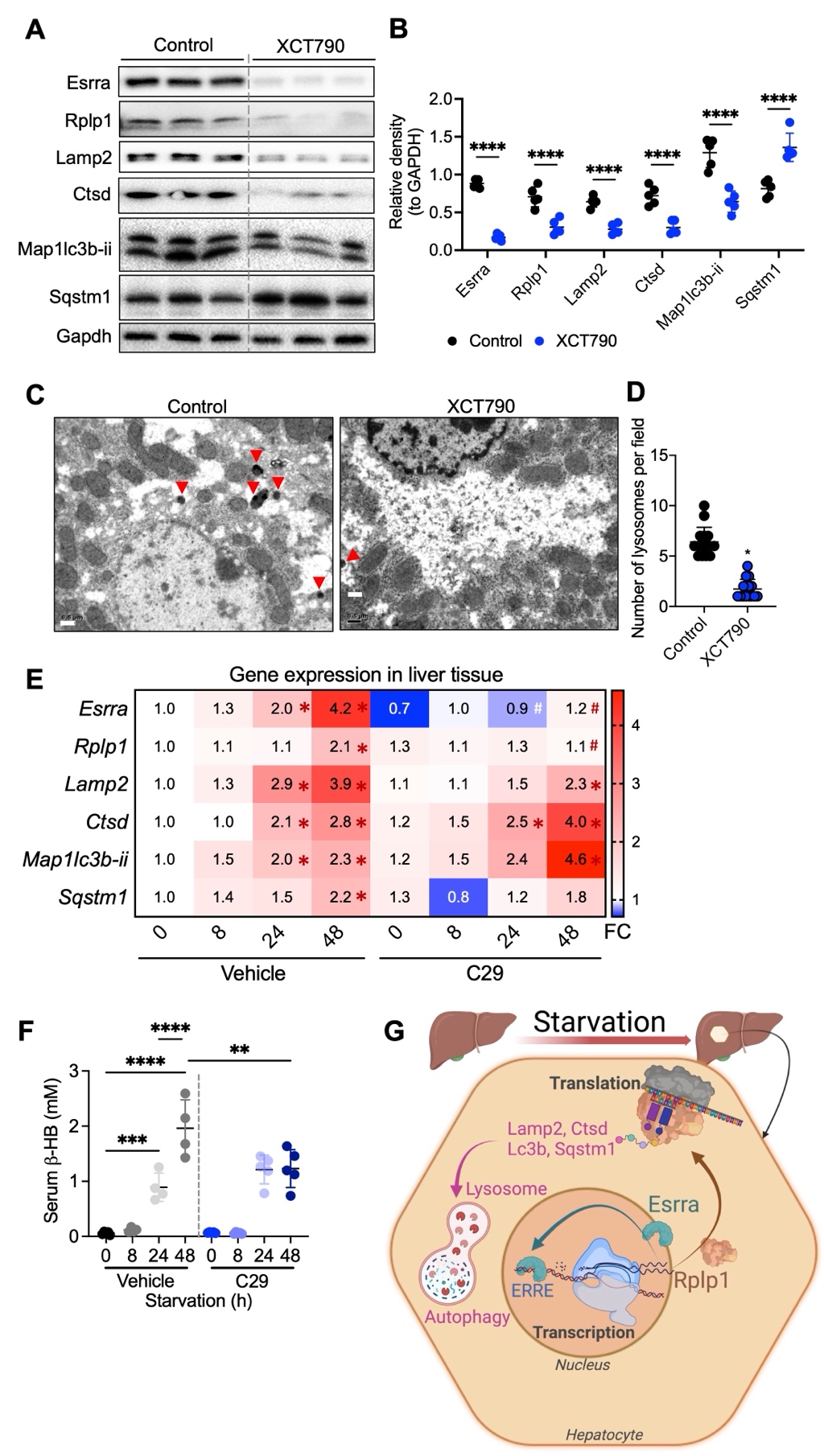

**Suppl. Fig. 4. Esrra regulated temporal changes in Rplp1-lysosome axis and autophagy during starvation *in vivo*.** (A) Representative Western blots of livers from vehicle control or XCT790 (a small molecule inhibitor of Esrra) treated mice (n=5 per groups). (B) Plots represent relative density of corresponding Western blots normalized to Gapdh. (C) Representative transmission electron micrograph of livers tissues from vehicle control or XCT790 treated mice (n=3 per group). Scale bar, 0.5 μm. (D) Lysosomes were counted and data are representative of 10 fields per group from three independent analyses. (E) RT-qPCR analysis of 0, 8, and 24 h starvation in mice livers at indicated time points treated with vehicle or Esrra-specific inverse agonist C29 (n=5 per group). (F) Serum β-hydroxybutyrate measurements in mice as described in panel E. (G) Illustration summarizing hepatic Esrra-Rplp1-lysosome axis regulation under starvation. Our findings show Esrra activation is required for upregulation of ribosomal protein Rplp1 during starvation that leads to translational recovery of lysosomal and autophagy proteins. Esrra transcriptionally regulate Rplp1 and its own expression during starvation followed by a translational recovery of lysosomal and autophagy proteins that is essential for successful recycling of macromolecules for selective biosynthesis of survival-related proteins during starvation.

Levels of significance: *P<0.05; **P<0.01; ***P<0.001; ****P<0.0001.

**Supplementary Methods**

*Animal studies*

**Fed-Fast-Refed mice:** 8-10 weeks old male C57BL/6J mice (n=5/group) were fed with normal chow control diet (NCD) and starved overnight before starting the experiment for synchronization. After one day, mice were kept in new empty cages (without husk bedding and NCD) for 24 (Fast group), and few mice from Fast group were refed for 6 h with NCD (Refed). Mice those were not undergone fasting or refeding after synchronization considered as Fed group.

**Starvation studies:** 8-10 weeks old male C57BL/6J mice (n=5/group) were fed with normal chow control diet (NCD) and starved overnight before starting the experiment for synchronization. After one day, mice were kept in new empty cages (without husk bedding) for 8, 24, 48 h. C29 (10 mg/kg body weight) was started injecting *i.p.* from the synchronization day to 48 h fasting day for the chronic inhibition of Esrra ^2^. Cycloheximide (Chx; 10 mg/kg body weight) was injected *i.p.* 60 min before euthanization for translation inhibition, while puromycin (Puro, 20 mg/kg body weight) was injected *i.p.* 30 min before euthanization for puromycin labeling of proteins to understand rate of translation.

***Esrra* KO mice:** *Esrra* KO mice are described elsewhere ^2, 3^. Esrra WT and *Esrra* KO mice (n=5/group) were synchronized by overnight fasting and starved for 24 h after two days or left fed. Later, mice were euthanized, blood and liver tissues were collected for mRNA and protein analysis.

***Alb*-m*ESRRA* overexpression:** For liver-specific expression, we used AAV8 mediated gene delivery of m*Esrra* gene cloned under the control of mouse *Alb* promoter (Vector Biolabs, USA). 8 weeks old male C57BL/6J mice (n-5/group) were used for liver-specific *Esrra* overexpression (*Alb-mEsrra*). Mice were generated by injecting AAV8-*Alb*-*mEsrra* (5X1011 gc/mice) via tail vein and housed for four weeks with no other intervention ^4^. We injected AAV8-*Alb*-Null which does not contain any DNA sequence under the transcriptional control of the *Alb* promoter as controls. Later, mice were euthanized, blood and liver tissues were collected for mRNA and protein analysis.

**General mouse care and ethics statement:** mice were purchased from InVivos, Singapore and, housed in hanging polycarbonate cages under a 12 h/12 h light/dark schedule at Duke-NUS vivarium. Mice were simple randomized before grouping and fed different diets and normal water, or fructose treated *ad libitum*. Animals were euthanized in CO2 chambers. All mice were maintained according to the Guide for the Care and Use of Laboratory Animals (NIH publication no. One.0.0. Revised 2011), and the experiments performed were approved by the IACUCs at SingHealth (2015/SHS/1104) and (2020/SHS/1549).

*Serum triglycerides and β-hydroxybutyrate (β-HB/Ketone bodies) measurements*

Serum triglycerides (TG), and β-HB was measured using Triglyceride Colorimetric Assay Kit (#10010303, Cayman) and β-HB (Ketone Body) Colorimetric Assay Kit (#700190, Cayman).

*Cell cultures*

**AML12 cells** (ATCC® CRL-2254™) were cultured as indicated elsewhere ^5, 6^. Starvation medium (DMEM:F12 mix with Pen/Strep lacking serum, ITS and Dexamethasone) was used for serum starvation experiments. To analyze the rate of translation, puromycin (10 μg/ml) was added for 15 min before harvest ^7^ whereas C29 (5 μM) was added to media for indicated time to inhibit Esrra ^2^. Bafilomycin A1 (5 nM) was used to analyze autophagy flux ^1^.

**Primary human hepatocytes** (5200, ScienCell) were cultured as indicated elsewhere ^4, 8^. Puromycin (10 μg/ml) was added for 15 min before harvest ^7^ whereas C29 (5 μM) was added to media for indicated time ^2^.

*Gene manipulation in cultured cells*

**Gene knockdown *in vitro*:** Silencer Select siRNAs (s4829, s4830, and s4831; Life Technologies Inc.) or ON-TARGETplus Smartpool siRNAs (L-040772-00-0010, Dharmacon) against *Esrra*, and Silencer Select siRNA (s234520; Life Technologies Inc.) for *Rplp1* gene knockdown were used in AML12 cells. Negative siRNA (Silencer Negative Control No. 1 siRNA; AM4611, Life Technologies Inc.) was used as a negative control. Transfections were carried out in AML12 cells in a 12-well or 6-well plate or four-well chambered slides or 24-well Seahorse XF plate using 30 nM of the above indicated siRNAs and negative control siRNA with Lipofectamine RNAiMAX (Invitrogen; Life Technologies Inc.) following the reverse transfection protocol, as indicated elsewhere ^1, 5^.

**Gene overexpression *in vitro*:** ORF sequence of m*Esrra* (Gene ID: 26379) and m*Rplp1* (Gene ID: 56040) genes were cloned in a pcDNA3.1(+)-C-6His plasmid by Genescript Limited (Hong Kong) to get Esrra_OMu13026C_pcDNA3.1(+)-C-6His and Rplp1_OMu11118C_pcDNA3.1(+)-C-6His constructs for Esrra and Rplp1 overexpression in AML12 cells respectively. Transfections were carried out in a 12-well plate or 6-well plate using Lipofectamine 3000 (Invitrogen; Life Technologies Inc.) following the reverse transfection protocol, as described elsewhere ^1^. Empty pcDNA3.1(+)-C-6His vector was used as control.

*RNA isolation and RT-qPCR analysis of gene expression*

RNA isolation and RT-qPCR for measuring gene expression were performed as described earlier ^1^. Predesigned KiCqStart SYBR Green optimized primers from Sigma-Aldrich (KSPQ12012) were used for RT-qPCR.

*Protein isolation and Western blotting analysis of protein expression*

Protein isolation and Western blotting analysis of protein expression were performed as described earlier ^1^. Primary antibodies for puromycin (1:25,000 dilution; MABE343, Merck), 1:1000 dilution of Esrra (ab16363, Abcam; 07-662, Millipore; and 13826S, CST), Rplp1 (PA5-103540, Invitrogen), Lamp2 (PA1-655, Invitrogen), Ctsd (sc377299, Santa Cruz), Gapdh (2118, CST), cleaved-Casp3 (9661, CST), Map1lc3b-ii/Lc3b (a2775, CST), and Sqstm1/p62 (5114, CST) were used. Horseradish peroxidase–conjugated secondary antibodies recognizing mouse (sc-2954) and rabbit (sc-2955) immunoglobulin Gs (IgGs) were purchased from Santa Cruz Biotechnology. Blots were observed on Image Lab software (BioRad) and densitometric analysis was performed using ImageJ software (NIH, Bethesda, MD, USA) normalized to GAPDH as loading controls.

*Immunoprecipitation of puromycin-incorporated proteins*

Protein immunoprecipitation was performed using Dynabeads™ Protein G for Immunoprecipitation (Invitrogen) as per manufacturer’s protocol. Non-denaturing lysis buffer (Abcam) was used to prepare tissue homogenate. Puromycin (1:5,000 dilution; MABE343, Merck), or 4 µg normal rabbit IgG as control (12-370, Sigma-Aldrich) was used for pull-down assay. Western blot analysis for the detection of pulled down proteins was performed as described above.

*Label-free quantitative proteomic analysis*

Label-free quantitative analysis of proteins, recovered from pooled triplicates each group, by LC-MS/MS was performed by NonovogeneAIT (Singapore) as described below.

Protein Quality Test: BSA standard protein solution was prepared according to the instructions of Bradford protein quantitative kit, with gradient concentration ranged from 0 to 0.5 g/L. BSA standard protein solutions and sample solutions with different dilution multiples were added into 96-well plate to fill up the volume to 20 µL, respectively. Each gradient was repeated three times. The plate was added 180 μL G250 dye solution quickly and placed at room temperature for 5 minutes, the absorbance at 595 nm was detected. The standard curve was drawn with the absorbance of standard protein solution and the protein concentration of the sample was calculated. 20 μg of the protein sample was loaded to 12% SDS-PAGE gel electrophoresis, wherein the concentrated gel was performed at 80 V for 20 min, and the separation gel was performed at 120 V for 90 min. The gel was stained by coomassie brilliant blue R-250 and decolored until the bands were visualized clearly.

Trypsin treatment: 120 μg of each protein sample was taken and the volume was made up to 100 μL with dissolution buffer, 1.5 μg trypsin and 500 μL of 100 mM TEAB buffer were added, sample was mixed and digested at 37 °C for 4 h. Andt hen,1.5 μg trypsin and CaCl2 were added, sample was digested overnight. Formic acid was mixed with digested sample, adjusted pH under 3, and centrifuged at 12000 g for 5 min at room temperature. The supernatant was slowly loaded to the C18 desalting column, washed with washing buffer (0.1% formic acid, 3% acetonitrile) 3 times, then eluted by some elution buffer (0.1% formic acid, 70% acetonitrile). The eluents of each sample were combined and lyophilized.

LC-MS/MS Analysis: Mobile phase A (100% water, 0.1% formic acid) and B solution (80% acetonitrile, 0.1% formic acid) were prepared. The lyophilized powder was dissolved in 10 μL of solution A, centrifuged at 14,000 g for 20 min at 4 °C, and 1 μg of the supernatant was injected into a home-made C18 Nano-Trap column (2 cm×75 μm, 3 μm). Peptides were separated in a home-made analytical column (15 cm×150 μm, 1.9 μm), using a linear gradient elution. The separated peptides were analyzed by Q Exactive HF-X mass spectrometer (Thermo Fisher), with ion source of Nanospray Flex™（ESI, spray voltage of 2.3 kV and ion transport capillary temperature of 320°C. Full scan range from m/z 350 to 1500 with resolution of 60000 (at m/z 200), an automatic gain control (AGC) target value was 3×106 and a maximum ion injection time was 20 ms. The top 40 precursors of the highest abundant in the full scan were selected and fragmented by higher energy collisional dissociation (HCD) and analyzed in MS/MS, where resolution was 15000 (at m/z200), the automatic gain control (AGC) target value was 1×105, the maximum ion injection time was 45 ms, a normalized collision energy was set as 27%, an intensity threshold was 2.2×10^4^, and the dynamic exclusion parameter was 20 s. The raw data of MS detection was named as “.raw”.

The identification and quantitation of protein: All resulting spectra were searched against Mus_musculus_uniprot_2019.01.18.fasta (85165 sequences) database by the search engines: Proteome Discoverer 2.2 (PD 2.2, Thermo). The search parameters are set as follows: mass tolerance for precursor ion was 10 ppm and mass tolerance for product ion was 0.02 Da. Carbamidomethyl was specified as fixed modifications, Oxidation of methionine (M) was specified as dynamic modification, and acetylation was specified as N-Terminal modification in PD 2.2. A maximum of 2 missed cleavage sites were allowed. In order to improve the quality of analysis results, the software PD 2.2 further filtered the retrieval results: Peptide Spectrum Matches (PSMs) with a credibility of more than 99% was identified PSMs. The identified protein contains at least 1 unique peptide. The identified PSMs and protein were retained and performed with FDR no more than 1.0%. The protein quantitation results were statistically analyzed by T-test. The proteins whose quantitation significantly different between experimental and control groups, (p < 0.05 and |log2FC| >= 2 (ratio >= 4 or ratio <= 0.25 [fold change, FC]), were defined as differentially expressed proteins (DEP).

The functional analysis of protein and DEP: Gene Ontology (GO) and KEGG (Kyoto Encyclopedia of Genes and Genomes) were used to analyze the protein family and pathway whereas Targeting Protein-Protein Interactions (PPIs) for protein-protein interactions on EnrichR platform (the Ma'ayan Lab, NY, USA) ^9-11^.

*Fluorescence imaging of the cells*

Autophagy flux analysis: ptfLC3 (Addgene plasmid No. 21074) was transfected using lipofectamine 3000 reagent (Invitrogen) in *Esrra* knockdown or control cells as described in manufacturer's protocol. ptfLC3 was a gift from Tamotsu Yoshimori and described elsewhere ^12^. After 48 h of transfection, cells were serum starved for further 24 h. The cells were then fixed with 4% paraformaldehyde for 15 min and washed three times with PBS. Slides were washed and wet mounted in VECTASHIELD Antifade Mounting Medium. Fluorescence imaging was performed using LSM710 Carl Zeiss (Carl Zeiss Microscopy GmbH, Oberkochen, Germany) confocal microscope at 40 × magnification.

Acridine orange (AO) staining: Cells were transfected and grown in 24-well plate for 48 h and serum starved for further 24 h. Thereafter, cells were incubated with 1 μg/ml of AO (Sigma-Aldrich) in PBS for 30 min at 37 °C, and observed under a fluorescence microscope as described elsewhere ^13^.

*Transmission electron microscopic (TEM) imaging of the liver tissues*

Fresh liver tissue was placed in fixative [2% paraformaldehyde and 3% glutaraldehyde in cacodylate buffer (pH 7.4)] and stored at 4°C. TEM analysis was performed as described previously ^1^. Images were taken using the Olympus EM208S transmission electron microscope (Japan) at ×10,000 magnifications. Mean number of lysosomes per TEM field from untreated control and XCT790-treated mouse liver samples was calculated from a total of 10 random fields per treatment.

*Mitochondrial oxygen consumption rate (OCR) measurement by Seahorse extracellular flux analyzer*

Seahorse extracellular flux analyser XFe96 (Agilent) was used for mitostress test, and mitochondrial fatty acid fuel oxidation analysis. Seahorse Wave Desktop software was used for report generation and data analysis while GraphPad PRISM 9 was used for statistical analysis and data presentation. 10,000 AML12 cells were seeded on XFe-96-well culture microplates and the assays were performed as described previously ^14^.

*Chromatin Immuno-Precipitation (ChIP)-qPCR in AML12 cells and ChIPseq in liver tissues*

ChIP-qPCR was performed as described previously ^1, 15^. For Esrra binding on the *Esrra* gene, primer pairs [ATGCATGGTCCCA- GAGTCAG (forward) and CTGGTTTGCGAGTTCCTCAA (reverse)] were used to amplify −629 to −498 region on *Esrra* promoters in AML12 cells. For Polr2a binding, a primer pair [ATTAGCATAGGGCACCTGGC (forward) and CGACCAC- CGTGGCTGAC (reverse)] was used to amplify −69 to +11 region in AML12 cells. However, on the *Rplp1* gene promoter, for Esrra binding, primer pairs [AGCTTCTTTGTGGTCCTGAGATT (forward) and ACCCCTCAGGTTAGGGTACA (reverse) were used to amplify −988 to −837 region on the *Rplp1* promoter in AML12 cells. For Polr2a binding on *Rplp1* gene promoter, a primer pair [CAATCGCACCGGAAGTCGAA (forward) and GACCAGTCCACCTATATACGCC (reverse)] were used to amplify −140 to +18 region in AML12 cells.

ChIPseq methodology is provided elsewhere ^2^.

*Polysome Profiling in vitro Using Sucrose Gradients*

AML12 cells were cultured with BSA (0.5% as control), PA (0.5 mM for 24 h) or PA with si*Esrra*, washed with cold PBS and subsequently lysed on ice using a lysis buffer containing Tris-Cl (pH 7.4, 20mM), NaCl (150mM), MgCl2 (5mM), NP40 (1%), DTT (1µM), Cycloheximide (100µg/ml), and cOmplete™ EDTA-free Protease Inhibitor (1X). The lysate was then homogenized by passing through a 27G needle five times. Cellular debris was separated by centrifuging at 20,000xg for 10 minutes at 4°C. Following this, the lysate was loaded onto a 10-50% sucrose gradient, which had been previously prepared with the Gradient Master. Polysome profiling was subsequently conducted, and fractions were collected using the Biocomp fractionator, in tandem with TRIAX. The data was plotted using GraphPad PRISM software.

*Statistical analysis*

Individual culture experiments were performed in triplicate and repeated at least three times independently using matched controls; the data were pooled (n=3/group), and statistical analysis was performed. Animal studies were performed as per the approved IACUC protocol, and the statistical analysis were performed (n=4-5/group) as per the biostatistician’s advice. Results are expressed as mean ± SD for all in vitro and in vivo experiments. Normality of the data was analyzed by Shapiro-Wilk test, then a parametric analysis was performed using unpaired student’s t-test, one-way ANOVA or two-way ANOVA followed by Tukey’s multiple-comparisons test, wherever applicable. The statistical significance of differences was assessed as *P<0.05; **P<0.01; ***P<0.001; ****P<0.0001). All statistical tests were performed using Prism 9 for Mac OS X (GraphPad Software).

**Tables**

Supplementary Table 1-8.

*Supplementary tables 1-8 can be found in one excel file submitted separately as Supplementary tables_proteomic pathways analysis.*
