## Supplemental tables for "Estrogen receptor-related receptor (Esrra) induces ribosomal protein Rplp1-mediated adaptive hepatic translation during prolonged starvation"

**Table 1: Venn analysis for KEGG pathways at 48 and 72 h serum starvation**

| 3 elements included exclusively in "48 h": | 15 common elements in "48 h" and "72 h": |
|---|---|
| **NAME** | **NAME** |
| Chemical carcinogenesis | Peroxisome |
| Drug metabolism | Thermogenesis |
| Proximal tubule bicarbonate reclamation | Mineral absorption |
| | Lysosome |
| | Fatty acid degradation |
| | PPAR signaling pathway |
| | Fatty acid elongation |
| | Valine, leucine and isoleucine degradation |
| | Other glycan degradation |
| | Oxidative phosphorylation |
| | Retrograde endocannabinoid signaling |
| | beta-Alanine metabolism |
| | Tryptophan metabolism |
| | SNARE interactions in vesicular transport |
| | Lysine degradation |

**19 elements included exclusively in "72 h":**

**NAME**

Ribosome
RNA transport
Propanoate metabolism
Huntington disease
Parkinson disease
Non-alcoholic fatty liver disease (NAFLD)
DNA replication
Alzheimer disease
Biosynthesis of unsaturated fatty acids
Primary bile acid biosynthesis
Endocytosis
RNA degradation
Amino sugar and nucleotide sugar metabolism
Glycosaminoglycan degradation
Glyoxylate and dicarboxylate metabolism
mRNA surveillance pathway
Ribosome biogenesis in eukaryotes
Spliceosome
Butanoate metabolism

#### Table 2: KEGG analysis for exclusive proteins at 72 h serum starvation

| Term | P-value | Adjusted P-value |
|---|---|---|
| **Ribosome** | **3.33E-19** | **7.47E-17** |
| **Lysosome** | **3.90E-08** | **4.37E-06** |
| **RNA transport** | **6.37E-07** | **4.76E-05** |
| **Thermogenesis** | **1.30E-06** | **7.27E-05** |
| DNA replication | 3.27E-06 | 1.47E-04 |
| RNA degradation | 1.37E-05 | 5.13E-04 |
| Huntington disease | 1.86E-05 | 5.94E-04 |
| **Oxidative phosphorylation** | **6.92E-05** | **0.001938846** |
| Alzheimer disease | 2.90E-04 | 0.007220182 |
| **Parkinson disease** | **5.36E-04** | **0.012011927** |
| Valine, leucine and isoleucine degradation | 7.34E-04 | 0.014949046 |
| Non-alcoholic fatty liver disease (NAFLD) | 8.19E-04 | 0.015278737 |
| Endocytosis | 0.0010629 | 0.018207378 |
| mRNA surveillance pathway | 0.00113796 | 0.018207378 |
| Propanoate metabolism | 0.0013275 | 0.019824003 |
| Amino sugar and nucleotide sugar metabolism | 0.00194922 | 0.027289148 |
| Retrograde endocannabinoid signaling | 0.00252494 | 0.032274563 |
| Mismatch repair | 0.00259349 | 0.032274563 |
| Small cell lung cancer | 0.00339165 | 0.0399858 |
| **Ribosome biogenesis in eukaryotes** | **0.00394515** | **0.044185689** |

**Genes**

**RPL4;RPL30;RPL34;RPLP1;MRPS12;RPLP0;RPL11;RPL8;RPL7;RPS15;RPS4X;RPL7A;RPS17;RPS16;**

**DNASE2A;CTSZ;HEXA;M6PR;AP3D1;GNS;GM2A;CTSL;LAMP2;PSAP;CTSH;TPP1;MAN2B1;AP3S1**

**EIF4A2;SEC13;PABPC4;RPP25L;NXF1;FXR2;EIF3G;NUP43;EIF3E;PABPC1;EIF3F;EIF4E2;EIF3C;EIF4**

**SMARCE1;NDUFA9;NDUFB9;SMARCC1;SMARCD2;NDUFA7;SMARCB1;SMARCC2;NDUFA11;NDU**

RFC5;RFC3;MCM7;RFC2;RPA1;MCM4;MCM6;MCM2

EDC4;DDX6;EXOSC6;CNOT1;PABPC4;XRN2;CNOT3;PABPC1;DCP1A;CNOT9;PFKP

NDUFA9;NDUFB9;NDUFA7;NDUFA11;NDUFB10;NDUFB5;HDAC1;DCTN1;DCTN4;NDUFB3;AP2A1;

**NDUFA9;NDUFB9;NDUFA7;NDUFA11;NDUFB10;NDUFB5;NDUFB3;SDHD;NDUFS8;UQCRFS1;ND**

NDUFA9;APP;NDUFB9;NDUFA7;NDUFA11;NDUFB10;LRP1;NDUFB5;NDUFB3;SDHD;NDUFS8;CAP

**NDUFA9;NDUFB9;NDUFA7;NDUFS8;NDUFA11;NDUFB10;NDUFB5;NDUFB3;UQCRFS1;NDUFS2;**

MCEE;PCCA;BCKDHB;MCCC1;ACADM;HADH;ACAT1

NDUFA9;NDUFB9;NDUFA7;NDUFS8;NDUFA11;NDUFB10;NDUFB5;NDUFB3;UQCRFS1;NDUFS2;S

ARFGEF1;WASHC5;ARFGEF2;RAB4A;GBF1;AGAP1;AP2A1;STAM;SNX4;HGS;KIF5B;CHMP4C;EPS15

NXF1;CPSF1;WDR82;PABPC4;CSTF3;CSTF2;CPSF2;PABPC1;WDR33

MCEE;PCCA;BCKDHB;ACADM;ACAT1

NAGK;GMDS;AMDHD2;HEXA;PGM3;RENBP

NDUFA9;NDUFB9;FAAH;NDUFA7;NDUFS8;NDUFA11;NDUFB10;NDUFB5;NDUFB3;NDUFS2;ND3

RFC5;RFC3;RFC2;RPA1

LAMA5;LAMB2;FN1;LAMC1;IKBKG;BAK1;NFKB1;PTK2

**NXF1;LSG1;CSNK2A1;XRN2;CSNK2B;SBDS;UTP14A;RPP25L;MDN1**

RPS15A;RPS19;RPL14;RPS3;RPL15;RPS11;RPS10;RPS13;RPL19;RPS9;RPL21;RPL22;RPL35A;RPL2[illegible]

[illegible]JFB10;NDUFB5;NDUFB3;SDHD;MTOR;SMARCA4;LIPE;NDUFS8;UQCRFS1;NDUFS2;SLC25A20;ND

7A;RPL37A;RPL24;RPS20;RPL22L1;RPS24

**Table 3: KEGG analysis for all upregulated proteins at 72 h serum starvation**

| Term | P-value | Adjusted P-value |
|---|---|---|
| **Ribosome** | **8.13E-16** | **2.46E-13** |
| **Lysosome** | **2.45E-13** | **3.72E-11** |
| **Peroxisome** | **4.34E-13** | **4.38E-11** |
| **Thermogenesis** | **5.66E-11** | **4.29E-09** |
| Valine, leucine and isoleucine degradation | 7.27E-09 | 4.41E-07 |
| **Fatty acid degradation** | **1.23E-08** | **6.22E-07** |
| **RNA transport** | **1.64E-07** | **7.09E-06** |
| **Oxidative phosphorylation** | **9.43E-07** | **3.57E-05** |
| Fatty acid elongation | 1.98E-06 | 6.65E-05 |
| Propanoate metabolism | 3.69E-06 | 1.12E-04 |
| Other glycan degradation | 5.07E-06 | 1.40E-04 |
| Huntington disease | 7.21E-06 | 1.82E-04 |
| Parkinson disease | 3.94E-05 | 9.18E-04 |
| PPAR signaling pathway | 5.71E-05 | 0.001236814 |
| Retrograde endocannabinoid signaling | 6.78E-05 | 0.001369709 |
| Non-alcoholic fatty liver disease (NAFLD) | 7.40E-05 | 0.001401306 |
| DNA replication | 8.58E-05 | 0.001528571 |
| Alzheimer disease | 1.66E-04 | 0.00279775 |
| Biosynthesis of unsaturated fatty acids | 3.19E-04 | 0.005090993 |
| Primary bile acid biosynthesis | 4.00E-04 | 0.006062109 |
| Mineral absorption | 4.61E-04 | 0.006650619 |
| Endocytosis | 5.07E-04 | 0.00697906 |
| RNA degradation | 7.41E-04 | 0.0097679 |
| Lysine degradation | 7.94E-04 | 0.010024894 |
| Tryptophan metabolism | 8.45E-04 | 0.010244404 |
| Amino sugar and nucleotide sugar metabolism | 9.73E-04 | 0.011341648 |
| Glycosaminoglycan degradation | 0.00156274 | 0.017537377 |
| Glyoxylate and dicarboxylate metabolism | 0.0016924 | 0.018314183 |
| beta-Alanine metabolism | 0.00200929 | 0.020993559 |
| **SNARE interactions in vesicular transport** | **0.00236902** | **0.023927054** |
| **mRNA surveillance pathway** | **0.00246538** | **0.024097114** |
| **Ribosome biogenesis in eukaryotes** | **0.00352191** | **0.03334812** |
| Spliceosome | 0.00403773 | 0.037073664 |
| Butanoate metabolism | 0.00502409 | 0.044773545 |

| Genes |
| --- |
| **RPL4;RPL30;RPL34;RPLP1;MRPS12;RPLP0;RPL11;RPL8;MRPL13;RPL7;RPS15;RPS4X;MRPL20;RP** |
| **SCARB2;CD63;HEXB;CTSZ;HEXA;LIPA;GNS;GM2A;CTSL;LAMP2;PSAP;NEU1;ATP6V0A2;CTSH;AP** |
| **PECR;ACOT8;PHYH;MVK;ACSL1;ECI2;ECH1;PIPOX;HSD17B4;CROT;GNPAT;NUDT7;AMACR;SCP2** |
| **NDUFB9;NDUFB8;SMARCD2;SMARCB1;NDUFA11;NDUFB10;NDUFB6;NDUFB11;NDUFB5;NDUF** |
| MCCC2;ACAA2;BCKDHB;MCCC1;ACAT1;HADHB;ALDH3A2;HADHA;MCEE;PCCA;EHHADH;PCCB;H |
| **ACADVL;CPT1A;ACAA2;ACSL1;ECI2;ACAT1;HADHB;ALDH3A2;HADHA;CPT2;EHHADH;ACADM;A** |
| **EIF4A2;EIF2B4;SEC13;SEH1L;NXT1;RPP30;PABPC4;THOC3;RPP25L;NXF1;FXR2;EIF3G;GEMIN5;N** |
| **NDUFA9;NDUFB9;NDUFB8;NDUFA7;NDUFA11;NDUFB10;NDUFB6;NDUFB11;NDUFB5;NDUFA1** |
| HADHB;HADHA;ACAA2;ACOT2;ACOT1;PPT2;HACD3;HADH;HACD2 |
| HADHA;MCEE;PCCA;EHHADH;BCKDHB;PCCB;ACADM;MLYCD;ACAT1 |
| MANBA;GLB1;MAN2B2;HEXB;HEXA;NEU1;MAN2B1 |
| NDUFA9;NDUFB9;NDUFB8;NDUFA7;NDUFA11;NDUFB10;NDUFB6;NDUFB11;NDUFB5;HDAC1;DC |
| NDUFA9;NDUFB9;NDUFB8;NDUFA7;NDUFA11;NDUFB10;NDUFB6;NDUFB11;NDUFB5;NDUFA10; |
| SLC27A1;CPT1A;ACSL1;APOA1;CPT2;SCP2;ACOX2;EHHADH;PLIN2;HMGCS2;ACADM;ACOX3;SLC2 |
| NDUFA9;NDUFB9;FAAH;NDUFB8;NDUFA7;NDUFA11;NDUFB10;NDUFB6;NDUFB11;NDUFB5;NDU |
| NDUFA9;NDUFB9;NDUFB8;NDUFA7;NDUFA11;NDUFB10;NDUFB6;NDUFB11;NDUFB5;NDUFA10; |
| RFC5;RFC3;MCM7;RFC2;RPA1;MCM4;MCM6;MCM2 |
| NDUFA9;APP;NDUFB9;NDUFB8;NDUFA7;NDUFA11;NDUFB10;LRP1;NDUFB6;NDUFB11;NDUFB5; |
| SCP2;ACOT2;HSD17B4;ACOT1;ACOX3;HACD3;HACD2 |
| ACOT8;AMACR;ACOX2;SCP2;HSD17B4 |
| FTL1;FTH1;HMOX1;SLC30A1;MT2;ATP1B1;STEAP2;SLC39A4 |
| ARFGEF1;WASHC5;ARFGEF2;RAB4A;IQSEC2;GBF1;SNX12;AGAP1;AP2A1;STAM;RAB22A;SNX4;IT |
| EDC4;DDX6;EXOSC6;CNOT1;PABPC4;XRN2;CNOT3;PABPC1;DCP1A;CNOT9;PFKP |
| ALDH3A2;HADHA;EHMT2;EHHADH;SETD1B;EHMT1;PIPOX;HADH;ACAT1 |
| ALDH3A2;HADHA;MAOB;EHHADH;CAT;KYAT3;HADH;ACAT1 |
| GNPDA1;NAGK;GMDS;HEXB;AMDHD2;HEXA;PGM3;RENBP |
| GLB1;HEXB;HEXA;GUSB;GNS |
| MCEE;PCCA;CAT;PCCB;ACO1;ACAT1 |
| ALDH3A2;HADHA;ALDH3B1;EHHADH;ACADM;MLYCD |
| **GOSR2;STX6;VAMP4;SNAP29;VAMP5;VTI1B** |
| **NXF1;CPSF7;NXT1;CPSF1;WDR82;PABPC4;CSTF3;CSTF2;CPSF2;PABPC1;WDR33** |
| **NXF1;LSG1;NXT1;CSNK2A1;RPP30;CSNK2A2;XRN2;CSNK2B;SBDS;UTP14A;RPP25L;MDN1** |
| SF3B4;HNRNPA3;DHX8;BUD31;DDX42;THOC3;PLRG1;CHERP;SYF2;SNRPE;SRSF5;CTNNBL1;HSPA1 |
| HADHA;EHHADH;HMGCS2;HADH;ACAT1 |

**.7A;RPS17;RPS16;RPS15A;RPS19;RPL14;RPS3;RPL15;RPS11;RPS10;RPS13;RPL19;RPS9;RPL21;RP**
**3S1;GUSB;CTSD;CTSC;CTSA;MANBA;DNASE2A;M6PR;AP3D1;GLB1;TPP1;MAN2B1;PPT2;GLA**

**A10;NDUFB3;LIPE;CPT2;UQCRFS1;SLC25A20;SMARCE1;NDUFA9;SMARCC1;CPT1A;NDUFA7;SMA**

**);NDUFB3;SDHD;NDUFS8;NDUFS5;ATP6V0A2;UQCRFS1;NDUFS2;ND1;ND3;ATP6V1C1**

TN1;DCTN4;NDUFA10;NDUFB3;AP2A1;SDHD;NRF1;NDUFS8;SIN3A;NDUFS5;POLR2C;UQCRFS1;NI

:H;HGS;KIF5B;CHMP4C;EPS15;RAB5A;HSPA1B;FGFR2;CYTH1;SNX6;SH3GL1;WASHC3

ARCC2;ACSL1;SDHD;ARID1B;MTOR;SMARCA4;NDUFS8;NDUFAF6;NDUFAF4;NDUFS5;NDUFAF2;I

NDUFS2;ND1;ND3;COX20

### Table 4: KEGG analysis for exclusively upregulated proteins at 48 h serum starvation

| Term | P-value | Adjusted P-value | Genes |
|---|---|---|---|

**Table 5: KEGG analysis for all upregulated proteins at 48 h serum starvation**

| **Term** | **P-value** | **Adjusted P-value** |
|---|---|---|
| Peroxisome | 3.86E-12 | 1.17E-09 |
| **Thermogenesis** | **2.55E-07** | **3.86E-05** |
| Mineral absorption | 4.38E-07 | 4.42E-05 |
| **Lysosome** | **6.96E-07** | **5.28E-05** |
| **Fatty acid degradation** | **1.38E-06** | **8.33E-05** |
| PPAR signaling pathway | 1.90E-05 | 9.58E-04 |
| Valine, leucine and isoleucine degradation | 3.14E-05 | 0.001360185 |
| Other glycan degradation | 3.60E-05 | 0.001361777 |
| Fatty acid elongation | 3.61E-05 | 0.001214754 |
| **Oxidative phosphorylation** | **2.07E-04** | **0.006261927** |
| Chemical carcinogenesis | 2.47E-04 | 0.006790431 |
| Retrograde endocannabinoid signaling | 5.45E-04 | 0.013773032 |
| Tryptophan metabolism | 6.53E-04 | 0.015210331 |
| beta-Alanine metabolism | 6.54E-04 | 0.014151683 |
| **SNARE interactions in vesicular transport** | **7.57E-04** | **0.015285403** |
| Drug metabolism | 0.00101785 | 0.01927546 |
| Proximal tubule bicarbonate reclamation | 0.00128674 | 0.022934226 |
| Lysine degradation | 0.00195181 | 0.032855492 |

**Genes**

PECR;PHYH;MVK;ACSL1;ECI2;PIPOX;HSD17B4;CROT;GNPAT;NUDT7;AMACR;ACOX2;EHHADH;CA[illegible]

**COA3;CPT1A;NDUFB8;NDUFB6;ACSL1;NDUFB11;COX15;NDUFA10;MAPK14;ARID1B;NDUFAF6;[illegible]**

FTL1;FTH1;HMOX1;MT2;ATP1B3;MT1;ATP1B1;STEAP2;SLC39A4

**SCARB2;CTSA;CD63;MANBA;ATP6AP1;HEXB;GBA;LIPA;AP3M1;GLB1;NEU1;ATP6V0A2;PPT2;CT[illegible]**

**HADHB;ALDH3A2;HADHA;CPT1A;ACAA2;CPT2;ACSL1;EHHADH;ECI2**

SLC27A1;CPT1A;CPT2;ACOX2;GK;ACSL1;EHHADH;DBI;HMGCS2;SORBS1

MCCC2;HADHB;ALDH3A2;HADHA;ACAA2;EHHADH;PCCB;HMGCS2

MANBA;GLB1;HEXB;GBA;NEU1

HADHB;HADHA;ACAA2;ACOT2;ACOT1;PPT2

**NDUFB8;ATP6AP1;NDUFB6;NDUFB11;COX15;NDUFS5;NDUFA10;ATP6V0A2;ND1;NDUFV3;ND5[illegible]**

CBR1;GSTM1;GSTA3;NAT2;ALDH3B1;MGST1;KYAT3;CYP3A13;UGT1A6B

NDUFB8;NDUFB6;NDUFB11;NDUFS5;NDUFA10;GNB1;ND1;NDUFV3;MAPK14;ND5;GNAI1

ALDH3A2;HADHA;MAOB;EHHADH;CAT;KYAT3

ALDH3A2;HADHA;ALDH3B1;EHHADH;MLYCD

**GOSR2;STX6;VAMP4;SNAP29;VAMP5**

GSTM1;TPMT;MAOB;NAT2;GSTA3;ALDH3B1;MGST1;UGT1A6B;NME1

GLUD1;ATP1B3;SLC25A10;ATP1B1

ALDH3A2;HADHA;SETD2;EHHADH;EHMT1;PIPOX

**CPT2;NDUFAF4;NDUFS5;NDUFAF2;ND1;NDUFV3;COA7;ND5;COX20**

### Table 6: Venn analysis for GO CC pathways at 48 and 72 h serum starvation

**17 elements included exclusively in "List 1":**

intrinsic component of mitochondrial inner membrane (GO:0031304)
integral component of mitochondrial inner membrane (GO:0031305)
signal recognition particle, endoplasmic reticulum targeting (GO:0005786)
preribosome (GO:0030684)
small-subunit processome (GO:0032040)
zonula adherens (GO:0005915)
integral component of mitochondrial outer membrane (GO:0031307)
intrinsic component of mitochondrial outer membrane (GO:0031306)
vacuole (GO:0005773)
signal recognition particle (GO:0048500)
myofibril (GO:0030016)
contractile fiber (GO:0043292)
proton-transporting V-type ATPase complex (GO:0033176)
mitochondrial inner membrane presequence translocase complex (GO:0005744)
mitochondrial intermembrane space (GO:0005758)
phagocytic vesicle (GO:0045335)
vacuolar proton-transporting V-type ATPase complex (GO:0016471)

**31 common elements in "List 1" and "List 2":**
mitochondrion (GO:0005739)
mitochondrial inner membrane (GO:0005743)
peroxisomal matrix (GO:0005782)
microbody lumen (GO:0031907)
peroxisomal part (GO:0044439)
microbody (GO:0042579)
peroxisome (GO:0005777)
azurophil granule (GO:0042582)
lysosome (GO:0005764)
lysosomal membrane (GO:0005765)
vacuolar lumen (GO:0005775)
mitochondrial outer membrane (GO:0005741)
lysosomal lumen (GO:0043202)
integral component of mitochondrial membrane (GO:0032592)
azurophil granule membrane (GO:0035577)
lytic vacuole (GO:0000323)
lytic vacuole membrane (GO:0098852)
secretory granule lumen (GO:0034774)
mitochondrial envelope (GO:0005740)
azurophil granule lumen (GO:0035578)
peroxisomal membrane (GO:0005778)
mitochondrial matrix (GO:0005759)
mitochondrial respiratory chain complex I (GO:0005747)
specific granule (GO:0042581)
Golgi subcompartment (GO:0098791)
late endosome (GO:0005770)
ficolin-1-rich granule lumen (GO:1904813)
late endosome membrane (GO:0031902)
primary lysosome (GO:0005766)
Golgi membrane (GO:0000139)
U2-type catalytic step 2 spliceosome (GO:0071007)

**46 elements included exclusively in "List 2":**
cytosolic ribosome (GO:0022626)
cytosolic part (GO:0044445)
focal adhesion (GO:0005925)
cytosolic large ribosomal subunit (GO:0022625)
large ribosomal subunit (GO:0015934)
cytosolic small ribosomal subunit (GO:0022627)
small ribosomal subunit (GO:0015935)
ribosome (GO:0005840)
nucleolus (GO:0005730)
cytoplasmic stress granule (GO:0010494)
cytoplasmic ribonucleoprotein granule (GO:0036464)
SWI/SNF complex (GO:0016514)
NuRD complex (GO:0016581)
CHD-type complex (GO:0090545)
BAF-type complex (GO:0090544)
vesicle coat (GO:0030120)
early endosome (GO:0005769)
COPI vesicle coat (GO:0030126)
COPI-coated vesicle membrane (GO:0030663)
npBAF complex (GO:0071564)
P-body (GO:0000932)
Set1C/COMPASS complex (GO:0048188)
nucleoplasm part (GO:0044451)
polysome (GO:0005844)
ficolin-1-rich granule (GO:0101002)
trans-Golgi network (GO:0005802)
mRNA cleavage and polyadenylation specificity factor complex (GO:0005847)
endoplasmic reticulum-Golgi intermediate compartment (GO:0005793)
cell cortex region (GO:0099738)
chromatin (GO:0000785)
polysomal ribosome (GO:0042788)
early endosome membrane (GO:0031901)
COPII-coated ER to Golgi transport vesicle (GO:0030134)
perinuclear region of cytoplasm (GO:0048471)
Sin3 complex (GO:0016580)
nuclear chromosome part (GO:0044454)
coated vesicle (GO:0030135)
autolysosome (GO:0044754)
Sin3-type complex (GO:0070822)
mitochondrial small ribosomal subunit (GO:0005763)
RNAi effector complex (GO:0031332)

RISC complex (GO:0016442)
stress fiber (GO:0001725)
contractile actin filament bundle (GO:0097517)
ribonucleoprotein granule (GO:0035770)
ER to Golgi transport vesicle membrane (GO:0012507)

### Table 7: GO CC analysis for all upregulated proteins at 72 h serum starvation

| Term | P-value |
|---|---|
| **mitochondrion (GO:0005739)** | **4.05E-24** |
| cytosolic ribosome (GO:0022626) | 1.17E-21 |
| mitochondrial inner membrane (GO:0005743) | 4.30E-19 |
| cytosolic part (GO:0044445) | 1.62E-18 |
| microbody lumen (GO:0031907) | 2.22E-14 |
| peroxisomal matrix (GO:0005782) | 2.22E-14 |
| peroxisomal part (GO:0044439) | 2.97E-13 |
| focal adhesion (GO:0005925) | 3.32E-13 |
| cytosolic large ribosomal subunit (GO:0022625) | 4.59E-13 |
| large ribosomal subunit (GO:0015934) | 1.17E-12 |
| peroxisome (GO:0005777) | 3.50E-12 |
| microbody (GO:0042579) | 3.50E-12 |
| **lysosome (GO:0005764)** | **1.63E-11** |
| lysosomal lumen (GO:0043202) | 4.94E-11 |
| cytosolic small ribosomal subunit (GO:0022627) | 8.54E-11 |
| small ribosomal subunit (GO:0015935) | 3.23E-10 |
| azurophil granule (GO:0042582) | 3.23E-10 |
| vacuolar lumen (GO:0005775) | 9.05E-10 |
| **ribosome (GO:0005840)** | **2.02E-09** |
| mitochondrial respiratory chain complex I (GO:0005747) | 1.63E-08 |
| **nucleolus (GO:0005730)** | **5.44E-08** |
| cytoplasmic stress granule (GO:0010494) | 7.92E-08 |
| secretory granule lumen (GO:0034774) | 8.07E-08 |
| cytoplasmic ribonucleoprotein granule (GO:0036464) | 2.30E-07 |
| lytic vacuole (GO:0000323) | 2.46E-07 |
| lysosomal membrane (GO:0005765) | 8.65E-07 |
| mitochondrial matrix (GO:0005759) | 1.05E-06 |
| SWI/SNF complex (GO:0016514) | 1.15E-06 |
| CHD-type complex (GO:0090545) | 1.97E-06 |
| NuRD complex (GO:0016581) | 1.97E-06 |
| BAF-type complex (GO:0090544) | 3.22E-06 |
| vesicle coat (GO:0030120) | 3.69E-06 |
| early endosome (GO:0005769) | 8.81E-06 |
| COPI vesicle coat (GO:0030126) | 2.06E-05 |
| azurophil granule lumen (GO:0035578) | 2.46E-05 |
| azurophil granule membrane (GO:0035577) | 2.74E-05 |
| COPI-coated vesicle membrane (GO:0030663) | 3.17E-05 |
| npBAF complex (GO:0071564) | 5.05E-05 |
| lytic vacuole membrane (GO:0098852) | 5.73E-05 |
| integral component of mitochondrial membrane (GO:0032592) | 1.08E-04 |
| P-body (GO:0000932) | 1.25E-04 |

| | |
|---|---|
| Set1C/COMPASS complex (GO:0048188) | 1.31E-04 |
| **polysome (GO:0005844)** | **2.96E-04** |
| nucleoplasm part (GO:0044451) | 3.00E-04 |
| ficolin-1-rich granule (GO:0101002) | 3.18E-04 |
| trans-Golgi network (GO:0005802) | 3.40E-04 |
| Golgi subcompartment (GO:0098791) | 3.43E-04 |
| mRNA cleavage and polyadenylation specificity factor complex (GO:0005847) | 5.47E-04 |
| Golgi membrane (GO:0000139) | 6.32E-04 |
| endoplasmic reticulum-Golgi intermediate compartment (GO:0005793) | 6.73E-04 |
| ficolin-1-rich granule lumen (GO:1904813) | 7.24E-04 |
| cell cortex region (GO:0099738) | 8.34E-04 |
| late endosome membrane (GO:0031902) | 8.45E-04 |
| chromatin (GO:0000785) | 8.72E-04 |
| polysomal ribosome (GO:0042788) | 9.65E-04 |
| early endosome membrane (GO:0031901) | 9.83E-04 |
| mitochondrial outer membrane (GO:0005741) | 0.00117922 |
| COPII-coated ER to Golgi transport vesicle (GO:0030134) | 0.001215952 |
| peroxisomal membrane (GO:0005778) | 0.001317085 |
| perinuclear region of cytoplasm (GO:0048471) | 0.001596977 |
| Sin3 complex (GO:0016580) | 0.001687867 |
| mitochondrial envelope (GO:0005740) | 0.002458954 |
| nuclear chromosome part (GO:0044454) | 0.002729172 |
| coated vesicle (GO:0030135) | 0.002998016 |
| **autolysosome (GO:0044754)** | **0.003623927** |
| Sin3-type complex (GO:0070822) | 0.00388167 |
| mitochondrial small ribosomal subunit (GO:0005763) | 0.005024094 |
| RISC complex (GO:0016442) | 0.005265498 |
| RNAi effector complex (GO:0031332) | 0.005265498 |
| contractile actin filament bundle (GO:0097517) | 0.006388383 |
| stress fiber (GO:0001725) | 0.006388383 |
| late endosome (GO:0005770) | 0.006616536 |
| ribonucleoprotein granule (GO:0035770) | 0.006669673 |
| U2-type catalytic step 2 spliceosome (GO:0071007) | 0.006892566 |
| primary lysosome (GO:0005766) | 0.007286878 |
| ER to Golgi transport vesicle membrane (GO:0012507) | 0.007503327 |
| specific granule (GO:0042581) | 0.008255963 |
| MLL3/4 complex (GO:0044666) | 0.009706799 |
| clathrin-coated vesicle (GO:0030136) | 0.009888073 |
| integral component of mitochondrial outer membrane (GO:0031307) | 0.010789579 |
| cytoplasmic dynein complex (GO:0005868) | 0.010789579 |
| Golgi-associated vesicle (GO:0005798) | 0.011018999 |
| histone acetyltransferase complex (GO:0000123) | 0.012753375 |
| intrinsic component of mitochondrial outer membrane (GO:0031306) | 0.012753375 |

| Adjusted P-value | Genes |
|---|---|
| **1.28E-21** | **MTCH1;SLC27A1;ACAA2;NDUFA11;ECI2;NDUFA10;CISD1;CLU;LACTB;CPNE3;MLY** |
| 1.86E-19 | RPL4;RPL30;RPL34;RPLP1;MRPS12;RPLP0;RPL11;RPL8;RPL7;RPS15;RPS4X;RPL7A;R |
| 4.54E-17 | NDUFA11;MRPS12;NDUFA10;CPT2;UQCRFS1;MPV17;ACAD11;SLC25A45;MRPL49; |
| 1.28E-16 | RPL4;RPL30;RPL34;RPLP1;MRPS12;RPLP0;RPL11;RPL8;RPL7;RPS15;RPS4X;RPL7A;R |
| 1.18E-12 | ACOT8;PHYH;ECI2;HSD17B4;PIPOX;CROT;GNPAT;NUDT7;AMACR;SCP2;ACOX2;EHH |
| 1.18E-12 | ACOT8;PHYH;ECI2;HSD17B4;PIPOX;CROT;GNPAT;NUDT7;AMACR;SCP2;ACOX2;EHH |
| 1.32E-11 | PECR;ACOT8;PHYH;ECI2;MGST1;PIPOX;HSD17B4;CROT;ALDH3A2;GNPAT;NUDT7;A |
| 1.32E-11 | RPL4;SCARB2;RPL30;RPLP1;RPLP0;FHL2;RPL8;RPL7;RPS15;RPS4X;RPL7A;RPS17;RPS |
| 1.62E-11 | RPL4;RPL30;RPL21;RPL34;RPLP1;RPLP0;RPL22;RPL11;RPL35A;RPL8;RPL7;RPL7A;FX |
| 3.70E-11 | RPL4;RPL30;RPL21;RPL34;RPLP1;RPLP0;RPL22;RPL11;RPL35A;RPL8;RPL7;RPL7A;FX |
| 9.25E-11 | PECR;PHYH;HSDL2;ECI2;ECH1;PIPOX;HSD17B4;CROT;GNPAT;NUDT7;AMACR;SCP2; |
| 9.25E-11 | PECR;PHYH;HSDL2;ECI2;ECH1;PIPOX;HSD17B4;CROT;GNPAT;NUDT7;AMACR;SCP2; |
| **3.97E-10** | **SCARB2;TINAGL1;CPQ;HEXB;HEXA;CTSZ;LIPA;GM2A;CTSL;ANPEP;LAMP2;ANXA6;** |
| 1.12E-09 | SCARB2;CHID1;CTSA;MANBA;HEXB;HEXA;LIPA;GNS;GM2A;CTSL;MAN2B2;GLB1;LA |
| 1.80E-09 | RPS9;MRPS12;RPS3A;RPS15;RPS4X;RPS17;RPS16;RPS15A;RPS19;RACK1;RPS3;RPS2 |
| 6.03E-09 | RPS9;MRPS12;RPS3A;RPS4X;RPS15;RPS17;RPS16;RPS15A;RPS19;RACK1;RPS3;RPS2 |
| 6.03E-09 | PYGB;CD63;RAB3D;HEXB;HEXA;MGST1;GNS;TTR;GM2A;CREG1;LAMP2;PSAP;TOM1 |
| 1.59E-08 | SCARB2;PYGB;HEXB;HEXA;LIPA;GNS;TTR;GM2A;CTSL;CREG1;LAMP2;PSAP;NEU1;G |
| **3.37E-08** | **RPL30;RPS9;MRPS12;RPL11;RRBP1;RPL8;MRPL13;RPS4X;RPL7A;RPS17;RPS19;RPS** |
| 2.59E-07 | NDUFA9;NDUFB9;NDUFB8;NDUFA7;NDUFA11;NDUFB6;NDUFB10;NDUFB11;NDUF |
| **8.21E-07** | **RPL4;EIF4A2;SRP19;ACADVL;SMARCB1;DDX42;CHD2;RPL7;SYNE1;PTBP1;RPL7A;R** |
| 1.11E-06 | GRB7;DDX6;MBNL1;MCRIP2;MCRIP1;PABPC4;CAPRIN1;IGF2BP1;CIRBP;PABPC1;EIF |
| 1.11E-06 | APP;PYGB;HEXB;PROS1;CTSZ;GNS;CLU;ACTR1B;TTR;GM2A;CREG1;NEU1;CTSH;GUS |
| 3.04E-06 | GRB7;DDX6;MBNL1;PABPC4;RPLP0;CIRBP;GIGYF2;SYNE1;EDC4;RPS4X;FXR2;MCRIP |
| 3.12E-06 | TINAGL1;CPQ;CTSZ;LIPA;CTSL;LAMP2;PSAP;NEU1;VPS11;CTSH;GUSB;CTSD;CTSC;V |
| 1.05E-05 | SCARB2;CD63;SEH1L;LRP1;RAB3D;MGST1;AP2A1;ANPEP;LAMP2;PSAP;VPS11;ATP6 |
| 1.24E-05 | LIPT2;ACADVL;NDUFB8;ACAA2;NAXE;ETFDH;IBA57;PDHB;ACAT1;MCEE;RPS3;MLY |
| 1.30E-05 | SMARCE1;SMARCC1;SMARCD2;SMARCC2;SMARCB1;ARID1B;SMARCA4 |
| 2.08E-05 | MBD3;CSNK2A1;HDAC1;APLP2;MTA2;GATAD2B;GATAD2A |
| 2.08E-05 | MBD3;CSNK2A1;HDAC1;APLP2;MTA2;GATAD2B;GATAD2A |
| 3.29E-05 | SMARCE1;SMARCC1;SMARCD2;SMARCC2;SMARCB1;ARID1B;SMARCA4 |
| 3.65E-05 | COPB2;COPA;SEC13;COPB1;COPG2;SEC24D;COPE;SEC31A;ARCN1 |
| 8.47E-05 | WASHC5;VAC14;LRP1;SNX12;PTPN23;RAB22A;RAB21;SNX4;VPS11;STX6;TOM1;WL |
| 1.92E-04 | COPB2;COPA;COPB1;COPG2;COPE;ARCN1 |
| 2.23E-04 | CTSA;PYGB;NHLRC3;GCA;HEXB;GNS;TTR;GM2A;GLB1;CREG1;MAN2B1;GUSB;GLA;C |
| 2.41E-04 | CD63;MANBA;TMEM30A;RAB3D;LAMP2;PSAP;TOM1;MGST1;CPNE3;DNAJC13;SNA |
| 2.72E-04 | COPB2;COPA;COPB1;COPG2;COPE;ARCN1 |
| 4.22E-04 | SMARCE1;SMARCC1;SMARCC2;SMARCB1;SMARCA4 |
| 4.65E-04 | SCARB2;STARD3;CD63;SEC13;SEH1L;LRP1;AP3D1;ATP11C;DNAJC13;MTOR;P2RX4; |
| 8.54E-04 | FUNDC2;APOO;CPT1A;SCO1;ABCB6;SCO2;ETFDH;BAK1;MICU1;TMEM70 |
| 9.69E-04 | EDC4;DDX6;CNOT1;CAPRIN1;AGO2;CNOT3;DCP1A;SQSTM1;EIF4E;CNOT9;SYNE1 |

| | |
|---|---|
| 9.90E-04 | RBBP5;WDR82;SETD1B;DPY30;ASH2L |
| **0.002160102** | **RPS4X;RPL7A;RPL30;FXR2;RPL11;AGO2;RPS3;RPL24;RPL8;RPL19** |
| 0.002160102 | ZCCHC8;GTF2A1;DHX8;GTF2B;CSTF2;EHMT1;PPWD1;DPY30;ASH2L;PTPN23;MTDH; |
| 0.002238207 | XRCC6;COPB1;CTSZ;UBR4;HUWE1;DYNLL1;GNS;DERA;DIAPH1;ACTR1B;GLB1;FTH1; |
| 0.002316649 | CHID1;ARFGEF1;APP;ARFGEF2;COG3;GBF1;M6PR;DPY30;ATP2C1;SCAMP2;RAB32; |
| 0.002316649 | SCARB2;COPB2;VAC14;COPA;SLC35B4;PROS1;COPB1;GOSR2;DPY30;ATP2C1;GALN |
| 0.003613643 | CPSF7;CPSF1;CPSF2;CSTF2;WDR33 |
| 0.004091223 | SCARB2;APP;COPB2;VAC14;COPA;SLC35B4;PROS1;COPB1;GOSR2;ATP2C1;GALNT1 |
| 0.004266091 | ERP44;MCFD2;SEC23IP;ANPEP;GOSR2;CTSZ;GBF1;FN1;STX5;COPG2;CTSC;NUCB2 |
| 0.004500295 | XRCC6;CTSZ;HUWE1;GNS;DERA;ACTR1B;GLB1;FTH1;CSNK2B;CAT;CTSH;GUSB;CTSD |
| 0.005055539 | ACTR1A;NUMA1;DCTN1;CTSZ |
| 0.005055539 | SCARB2;VAC14;CD63;STARD3;GOSR2;LAMP2;ANXA6;VTI1B |
| 0.005116195 | ANKRD17;SMARCD2;SMARCB1;MCM7;HDAC1;EHMT2;CTR9;PDS5A;SUPT6H;LOXL2 |
| 0.005561568 | RPL7A;RPL30;RPL11;RPL24;RPL8;RPL19 |
| 0.005566304 | RAB21;VAC14;SNX4;RAB4A;ANXA1;DNAJC13;RAB5A;WLS;VTI1B;SNX6 |
| 0.006558119 | CPT1A;MAOB;ABCB6;ACSL1;CISD2;RMDN3;NIPSNAP2;RAB32;FAM210B;HADHB;GP |
| 0.006645805 | MCFD2;SEC13;SEC23IP;GOSR2;CTSZ;STX5;SEC24D;CTSC;VTI1B;SEC31A |
| 0.007076544 | ALDH3A2;PECR;GNPAT;CAT;MGST1;HSD17B4;DECR2 |
| 0.008437364 | EIF4A2;APP;SET;GDPD1;CLU;MTDH;CDH1;KIF5B;VPS53;S100A13;RACK1;STX6;CAPN |
| 0.008771375 | MORF4L1;CSNK2A1;HDAC1;SIN3A |
| 0.012572392 | HADHB;MAOB;ABCB6;UQCC2;ABCB8;SDHD;SPG7;TIMM10;MICU1 |
| 0.013732501 | SMARCD2;SMARCB1;MCM7;TRRAP;HDAC1;EHMT2;CENPA;YY1;PURA;SFR1;RAD21; |
| 0.014849547 | SEC23IP;SNX18;CTSZ;NUMB;STX6;SNX9;CTSC;SEC31A;ARCN1 |
| **0.017673614** | **FTH1;LAMP2;SQSTM1** |
| 0.01864378 | MORF4L1;CSNK2A1;HDAC1;SIN3A |
| 0.023770714 | MRPS35;MRPS36;MRPS12;MRPS21;DAP3 |
| 0.024190764 | DDX6;AGO2;EIF4E |
| 0.024190764 | DDX6;AGO2;EIF4E |
| 0.028522782 | LIMA1;FBLIM1;FHOD1;PXN;PTK2;MYLK |
| 0.028522782 | LIMA1;FBLIM1;FHOD1;PXN;PTK2;MYLK |
| 0.028962826 | SCARB2;CHID1;STARD3;VAC14;CD63;GOSR2;PIK3R4;LAMP2;VPS11;ANXA6;VAMP5; |
| 0.028962826 | EDC4;RPS4X;DDX6;FXR2;RPLP0;CTSH;PABPC1;DCP1A;EIF4E |
| 0.029526263 | DHX8;SYF2;BUD31;SNRPE;PLRG1 |
| 0.030799206 | HEXB;HEXA;ANXA11 |
| 0.031296774 | MCFD2;SEC13;GOSR2;STX5;SEC24D;VTI1B;SEC31A |
| 0.033988835 | TMEM30A;CTSZ;UBR4;MOSPD2;ANXA11;ARMC8;NFKB1;ERP44;ALDH3B1;NEU1;TC |
| 0.039449428 | RBBP5;DPY30;ASH2L |
| 0.039677456 | SCARB2;SNX18;M6PR;NUMB;STX6;SNX9;AP2A1;VAMP4;STEAP2;RAB5A |
| 0.042225883 | FUNDC2;CPT1A;ABCB6;BAK1 |
| 0.042225883 | SNX4;DCTN4;DYNLL1;DYNLL2 |
| 0.042597839 | SEC23IP;COPB1;CTSZ;STEAP2;CTSC;SEC31A;ARCN1 |
| 0.048128809 | MORF4L1;ELP1;ELP2;ELP3 |
| 0.048128809 | FUNDC2;CPT1A;ABCB6;BAK1 |

**CD;MPV17;HADH;RARS2;MCCC2;CPT1A;ENDOG;ACSL1;ECH1;MCCC1;SDHD;BCS1L;GCAT;RAB32;**
PS17;RPS16;FXR2;RPS15A;RPS19;RACK1;RPL14;RPS3;RPL15;RPS11;RPS10;RPS13;RPL19;RPS9;RPL
MRPS21;TIMM22;SDHD;GCAT;HADHB;SLC25A16;MRPL53;HADHA;MRPL50;NDUFS5;SLC25A51;SL
PS17;RPS16;FXR2;RPS15A;RPS19;RACK1;RPL14;RPS3;IKBKG;RPL15;RPS11;RPS10;RPS13;RPL19;RP

16;RPS19;ANXA6;CAPN2;CPNE3;RPS11;RPS10;RPS13;TNS2;RPS9;ANXA1;RPL22;ITGA1;RPS3A;RPL

**CTSH;ENPP1;GUSB;CTSD;CTSC;CHID1;STARD3;SEC13;MANBA;ATP11C;HGS;VAMP4;PCYOX1;CD**

;CPNE3;GUSB;SNAP29;CTSC;CTSA;MANBA;TMEM30A;NHLRC3;GCA;ANXA11;DNAJC13;GLB1;MA
JSB;CTSD;CTSC;CHID1;CTSA;MANBA;NHLRC3;GCA;MAN2B2;GLB1;TPP1;MAN2B1;PPT2;GLA

**PS19;CPT2;CPNE3;RPS11;RPS10;UTP14A;RPS13;RPS9;NSUN2;SLC30A5;CIRBP;RPS3A;DDX52;GP**

B;CTSD;CTSC;VTI1B;CTSA;XRCC6;NHLRC3;GCA;XRCC5;FN1;HUWE1;APOA1;ARMC8;NFKB1;DERA;
2;CNOT1;MCRIP1;CAPRIN1;CNOT3;AGO2;IGF2BP1;CTSH;PABPC1;DCP1A;SQSTM1;CNOT9;EIF4E

V0A2;ANXA6;TOM1;ENPP1;CPNE3;SLC15A4;SNAP29;ATP6V1C1;VTI1B;VPS16;STARD3;SEC13;MA
D;ACADM;HMGCS2;HADH;DECR1;MCCC2;RARS2;NDUFA9;UQCC2;MCCC1;BCKDHB;NMNAT3;LA

S;VTI1B;VPS16;SNX6;RAB4A;ANXA1;VCAM1;APOA1;DNAJC13;RAB32;ITCH;HGS;STEAP2;RAB5A

ANPEP;LAMP2;PSAP;GNB1;VPS11;ATP6V0A2;ANXA6;ENPP1;LAMTOR3;SLC15A4;ATP6V1C1;VTI1B

;MED14;RBBP5;POLR2C;NUDT19;SBF1;USP7;UQCC2;SETD1B;WDR18;ELP1;ELP2;ELP3;GTF2F2;MC

T10;RAB21;APOO;CDH1;MAN2A1;STX6;STX5;SNX9;SNAP29;CYTH1;WLS;CHID1;RHBDF1;ARFGEF1;

);RAB21;APOO;MAN2A1;VPS53;STX6;STX5;SNAP29;CYTH1;WLS;RHBDF1;ARFGEF1;SLC30A6;ARFG

;YY1;SIN3A;POGZ;DPF2;MTA2;MBD3;SMARCE1;SMARCC1;SMARCC2;BUD31;GATAD2B;SMARCA4

N2;TLK2;HMOX1;DNM1L;EIF4E;VTI1B;SEC31A;TPD52;ARFGEF1;RAB4A;ANXA4;M6PR;CHERP;P2RX

POGZ;DPF2;MTA2;MBD3;SMARCE1;SMARCC1;XRCC6;SMARCC2;XRCC5;BUD31;RPA1;THOC3;GAT

**LARS2;ILF3;SCO1;SCO2;CARD19;TMEM126A;ACOT2;GUF1;DAP3;LIPT2;ABCB6;MAOB;NDUFB11**

C25A10;MICU1;SMIM20;DAP3;SLC25A13;NDUFB9;NDUFB8;MRPS35;NDUFB10;NDUFB6;MRPS36
S9;RPL21;RPL22;RPL35A;RPS3A;RPL27A;RPL37A;RPL24;GEMIN5;RPS20;RPL22L1;RPS24

37A;CAT;NUMB;PABPC1;GRB7;LRP1;FBLIM1;PXN;HACD3;SLC9A1;SLC9A3R2;RAB21;LIMA1;RPS3;

**63;SEH1L;LRP1;GNS;PSAP;NEU1;VPS11;ATP6V0A2;SLC15A4;ATP6V1C1;VTI1B;VPS16;CTSA;AP3D**

**RC5B;ILF3;KIF2A;WDR82;SUB1;GRWD1;XRN2;FRG1;SBDS;SRSF5;GTF3C1;DDX6;RPL11;MRPS31;**

;SLC30A6;ARFGEF2;COG3;GBF1;M6PR;ARCN1;RAB32;TPST2;RAB12;COPG2;MAN1B1;VAMP4;COP

**;MGST1;ABCB8;HACD3;ACAT1;PGS1;MRPL20;APOO;PRODH;TRMT1L;MARK2;DECR1;DNAJC19;**

;NDUFB11;NDUFB5;NDUFB3;MRPS31;TIMM10;SPG7;MRPL13;MRPL57;MRPL55;PGS1;MRPL20;T

**1;DNAJC13;MTOR;P2RX4;MAN2B2;GLB1;RAB12;GNB1;TPP1;MAN2B1;PPT2;LAMTOR3;GLA**

**ABCB8;NOL9;EXOSC6;RBBP5;S100A13;RPS3;LYAR;MDN1;ARFGEF1;XRCC6;NOP16;NIFK;KRR1;I**

**UQCC2;RMDN3;NFKB1;GLUD1;AMACR;NDUFAF6;NDUFAF4;NDUFAF2;ECHDC2;ACO1;COX20;TR**

MEM65;RPS3;RDH13;SLC25A20;PRODH;DNAJC19;NDUFA7;UQCC2;SQOR;EXOG;COQ7;NDUFAF6;N

MU;ACADVL;FUNDC2;CPT2;OPA3;RACK1;UQCRFS1;ACADM;HMGCS2;DNM1L;HSDL2;BCKDHB;

GLRX2;NIPSNAP1;NIPSNAP2;SLC25A16;HADHB;HADHA;NDUFS8;PLSCR3;NDUFS5;PCCB;CAT;RPU

ISD3;NDUFS2;SLC25A10;MICU1;FXN;SLC25A13;KANK2;NDUFB9;NDUFB8;GTF3C4;NDUFB6;NAXI

E;NDUFB5;MRPS36;MRPS31;IBA57;TIMM10;PDHB;SPG7;TMEM70;AKAP1;PDF;HS1BP3;SLC25A2

20;BAK1;NDUFA9;NDUFA7;NMNAT3;EXOG;PNKD;XPNPEP3;SARDH;COQ8B;SLIRP;CRAT;HSPA1B

#### Table 8: GO CC analysis for all upregulated proteins at 48 h serum starvation

| Term | P-value |
|---|---|
| **mitochondrion (GO:0005739)** | **6.63E-22** |
| mitochondrial inner membrane (GO:0005743) | 5.34E-14 |
| microbody lumen (GO:0031907) | 3.91E-12 |
| peroxisomal matrix (GO:0005782) | 3.91E-12 |
| peroxisomal part (GO:0044439) | 7.09E-12 |
| peroxisome (GO:0005777) | 1.85E-10 |
| microbody (GO:0042579) | 1.85E-10 |
| azurophil granule (GO:0042582) | 1.04E-08 |
| **lysosome (GO:0005764)** | **8.92E-08** |
| lysosomal membrane (GO:0005765) | 1.70E-07 |
| vacuolar lumen (GO:0005775) | 6.50E-07 |
| mitochondrial outer membrane (GO:0005741) | 1.71E-06 |
| lysosomal lumen (GO:0043202) | 2.10E-05 |
| integral component of mitochondrial membrane (GO:0032592) | 3.14E-05 |
| azurophil granule membrane (GO:0035577) | 4.08E-05 |
| lytic vacuole (GO:0000323) | 6.35E-05 |
| lytic vacuole membrane (GO:0098852) | 7.15E-05 |
| secretory granule lumen (GO:0034774) | 9.73E-05 |
| mitochondrial envelope (GO:0005740) | 1.44E-04 |
| azurophil granule lumen (GO:0035578) | 1.77E-04 |
| peroxisomal membrane (GO:0005778) | 2.37E-04 |
| mitochondrial matrix (GO:0005759) | 5.78E-04 |
| mitochondrial respiratory chain complex I (GO:0005747) | 9.06E-04 |
| intrinsic component of mitochondrial inner membrane (GO:0031304) | 0.003695644 |

| Adjusted P-value | Genes |
|---|---|
| **1.59E-19** | **TRMU;MTCH1;SLC27A1;ACAA2;ECI2;NDUFA10;CISD1;CLU;FUNDC2;CPT2;AIFM2;** |
| 6.38E-12 | NDUFB8;MRPS35;NDUFB6;NDUFB11;COX15;NDUFA10;TIMM10;SPG7;MRPL13;MR |
| 2.33E-10 | PHYH;ECI2;HSD17B4;PIPOX;CROT;GNPAT;NUDT7;AMACR;ACOX2;EHHADH;CAT;NU |
| 2.33E-10 | PHYH;ECI2;HSD17B4;PIPOX;CROT;GNPAT;NUDT7;AMACR;ACOX2;EHHADH;CAT;NU |
| 3.39E-10 | PECR;PHYH;ECI2;MGST1;PIPOX;HSD17B4;CROT;ALDH3A2;GNPAT;NUDT7;AMACR; |
| 6.31E-09 | PECR;PHYH;ECI2;PIPOX;HSD17B4;CROT;GNPAT;NUDT7;AMACR;ACOX2;ZADH2;EH |
| 6.31E-09 | PECR;PHYH;ECI2;PIPOX;HSD17B4;CROT;GNPAT;NUDT7;AMACR;ACOX2;ZADH2;EH |
| 3.11E-07 | CTSA;PYGB;CD63;MANBA;NHLRC3;GCA;RAB3D;HEXB;PRKCD;MGST1;ANXA11;BST2 |
| **2.37E-06** | **SCARB2;CD63;SEH1L;SLC44A2;HEXB;GBA;LIPA;GNAI1;AP3M1;FYCO1;ANPEP;NEU** |
| 4.06E-06 | SCARB2;STARD3;CD63;MANBA;SEH1L;SLC44A2;RAB3D;GBA;MGST1;ATP11C;GNAI1 |
| 1.41E-05 | SCARB2;CHID1;CTSA;PYGB;MANBA;NHLRC3;GCA;HEXB;PRKCD;GBA;LIPA;TTR;GLB1 |
| 3.41E-05 | CPT1A;MAOB;ABCB6;ACSL1;VPS13C;RAB32;FAM210B;HADHB;HAX1;MSTO1;GPA |
| 3.87E-04 | SCARB2;CHID1;CTSA;MANBA;GLB1;HEXB;GBA;NEU1;PPT2;LIPA |
| 5.36E-04 | FUNDC2;APOO;COA3;CPT1A;SCO1;ABCB6;SCO2;TMEM70 |
| 6.50E-04 | BST2;CD63;MANBA;VNN1;RAB3D;TOM1;MGST1;SNAP29 |
| 9.48E-04 | CHID1;CTSA;MANBA;USP4;GBA;LIPA;FYCO1;AP3M1;RAB12;NEU1;PPT2;VAMP4;CT |
| 0.001004524 | SCARB2;STARD3;CD63;SEH1L;SLC44A2;GBA;ATP11C;GNAI1;AP3M1;P2RX4;ANPEP; |
| 0.001292143 | CTSA;PYGB;NHLRC3;GCA;HEXB;PROS1;PRKCD;MAPK14;CLU;DERA;TTR;GLB1;NEU1; |
| 0.001808573 | HADHB;HAX1;MAOB;ABCB6;UQCC2;MICU2;SPG7;TIMM10 |
| 0.002114797 | CTSA;PYGB;TTR;NHLRC3;GCA;GLB1;HEXB;PRKCD;CTSC |
| 0.002698445 | ALDH3A2;PECR;GNPAT;CAT;MGST1;HSD17B4 |
| 0.006280193 | MCCC2;LIPT2;GADD45GIP1;NDUFB8;ACAA2;NAXE;UQCC2;PAM16;IBA57;LARS2;GL |
| 0.009409733 | NDUFB8;NDUFB6;NDUFB11;NDUFS5;NDUFA10;NDUFV3 |
| 0.036802459 | APOO;COA3;SCO1;SCO2 |

**MPV17;MLYCD;HMGCS2;COA7;MCCC2;CPT1A;ACSL1;NGDN;GLRX2;BCS1L;HADHB;RAB32;SLC25**

RPL57;PARL;PGS1;MRPL20;CPT2;MPV17;RDH13;NDUFV3;ACAD11;TIMMDC1;UQCC2;SQOR;MRPS

**1;ATP6V0A2;ENPP1;CTSC;CHID1;CTSA;STARD3;MANBA;USP4;ATP11C;P2RX4;GLB1;RAB12;GNB**

1;BST2;AP3M1;VNN1;P2RX4;ANPEP;GNB1;ATP6V0A2;TOM1;ENPP1;LAMTOR3;SNAP29;SIDT2

**A16;LARS2;HADHA;HAX1;SCO1;PLSCR3;SCO2;CARD19;NDUFS5;PCCB;RPUSD3;CAT;ACOT2;TMEI**
21;COQ7;HADHB;SLC25A16;HADHA;MRPL50;NDUFAF6;NDUFAF4;NDUFS5;NDUFAF2;SLC25A10;M

**M126A;GUF1;SLC25A10;MICU2;LIPT2;NDUFB8;ABCB6;NDUFB6;MAOB;NAXE;COX15;NDUFB11;[illegible]**

MGST1;IBA57;TIMM10;SPG7;TMEM70;AKAP1;PGS1;MRPL20;APOO;PDF;PDPR;PTPMT1;NDUFV

**3;GADD45GIP1;COA3;TIMMDC1;GK;UQCC2;GFER;BRI3BP;GLUD1;AMACR;ARMC10;NDUFAF6;N**

**DUFAF4;NDUFAF2;SLIRP;HSPA1B;BCL2L1;COX20**
